# Jekyll or Hyde? The genome (and more) of *Nesidiocoris tenuis*, a zoophytophagous predatory bug that is both a biological control agent and a pest

**DOI:** 10.1101/2020.02.27.967943

**Authors:** K. B. Ferguson, S. Visser, M. Dalíková, I. Provazníková, A. Urbaneja, M. Pérez-Hedo, F. Marec, J. H. Werren, B. J. Zwaan, B. A. Pannebakker, E. C. Verhulst

## Abstract

*Nesidiocoris tenuis* (Reuter) is an efficient predatory biological control agent used throughout the Mediterranean Basin in tomato crops but regarded as a pest in northern European countries. Belonging to the family Miridae, it is an economically important insect yet very little is known in terms of genetic information – no published genome, population studies, or RNA transcripts. It is a relatively small and long-lived diploid insect, characteristics that complicate genome sequencing. Here, we circumvent these issues by using a linked-read sequencing strategy on a single female *N. tenuis*. From this, we assembled the 355 Mbp genome and delivered an *ab initio*, homology-based, and evidence-based annotation. Along the way, the bacterial “contamination” was removed from the assembly, which also revealed potential symbionts. Additionally, bacterial lateral gene transfer (LGT) candidates were detected in the *N. tenuis* genome. The complete gene set is composed of 24,688 genes; the associated proteins were compared to other hemipterans (*Cimex lectularis*, *Halyomorpha halys*, and *Acyrthosiphon pisum*), resulting in an initial assessment of unique and shared protein clusters. We visualised the genome using various cytogenetic techniques, such as karyotyping, CGH and GISH, indicating a karyotype of 2n=32 with a male-heterogametic XX/XY system. Additional analyses include the localization of 18S rDNA and unique satellite probes via FISH techniques. Finally, population genomics via pooled sequencing further showed the utility of this genome. This is one of the first mirid genomes to be released and the first of a mirid biological control agent, representing a step forward in integrating genome sequencing strategies with biological control research.

## Introduction

Hemiptera is the fifth largest insect order and the most speciose hemimetabolous order with over 82,000 described species (Panfilio and Angelini, 2018). While recent sequencing projects have presented a variety of information about hemipteran genomes, large families such as the plant bugs Miridae still lack genomic resources, with the exception of transcriptomic resources for some members (Tian et al., 2015), and the more recent genome of *Apolygus lucorum*, a mirid pest that has a publicly available genome as of December 2019 (NCBI BioProject PRJNA526332). With the exception of *A. lucorum*, the lack of genomic resources for Miridae is in spite of the diverse life histories present, as it contains not only some of the most notorious agricultural pests but also predators that are often used in biological control (van Lenteren et al., 2018). In addition, Hemiptera are known for their intriguing karyotype evolution involving holocentric (holokinetic) chromosomes but there is a lack of cytogenetic information on Miridae. The absence of the ancestral TTAGG_n_ telomeric repeat have been reported for mirids *Macrolophus* spp.*, Deraeocoris* spp., and *Megaloceroea recticornis* (Geoffroy) (Grozeva et al., 2019, 2011; Jauset et al., 2015) but more knowledge of this trait is necessary for evolutionary studies of genomes and karyotypes. Furthermore, the taxonomic issues that lie within both Miridae and Hemiptera could better be resolved using protein and transcriptome-based analysis, but there is a noted lack of data in this regard a well (Panfilio and Angelini, 2018). While there is a relatively large amount of research into mirids and their use in biological control compared to other predators (Puentes et al., 2018), sequencing projects, if any, often focus on pest species and not on biological control agents (Panfilio and Angelini, 2018). For more advanced molecular methods such as RNAi and CRISPR-based genome editing strategies, it is necessary to have access to genomic and transcriptomic resources of the target species, and so these methods are currently out of reach for *N. tenuis* researchers. This lack in resources on both agricultural pests and biological control agents in the Miridae prompted us to generate genomic and cytogenetic resources of a mirid species that is both.

*Nesidiocoris tenuis* (Reuter) (Hemiptera: Miridae) is a zoophytophagous mirid used as a biological control agent worldwide, including in Spain, the Mediterranean Basin, and China (Pérez-Hedo and Urbaneja, 2016; Xun et al., 2016). Throughout the Mediterranean Basin, *N. tenuis* is used in tomato greenhouses and open fields against whiteflies (Hemiptera: Aleyrodidae), and the South American tomato pinworm, *Tuta absoluta* (Meyrick) (Lepidoptera: Gelechiidae) (Calvo et al., 2009; Mollá et al., 2014). In addition, due to its high degree of polyphagous behaviour, it is able to prey on other pest species such as thrips, leaf miners, leafhoppers, aphids, spider mites, and lepidopteran pests (Pérez-Hedo and Urbaneja, 2016). While *N. tenuis* is an important biological control agent in Mediterranean countries (Urbaneja et al., 2012), it is often cited as a pest in other contexts and countries (Calvo et al., 2009; Pérez-Hedo and Urbaneja, 2016). When prey is scarce in tomatoes, due to its phytophagy, *N. tenuis* can cause plant lesions such as brown discolouration around tender stems, known as necrotic rings, in addition to leaf wilt, and flower abortion (Arnó et al., 2010). This switch to phytophagy has been observed to be inversely proportional to the availability of prey (Sanchez, 2009). Therefore, much of the research thus far has focused on characterizing *N. tenuis* biology and ecology, classifying the induced damage, and attempting to reduce it (Biondi et al., 2015; Castañé et al., 2011; Garantonakis et al., 2018; Martínez-García et al., 2016; Urbaneja-Bernat et al., 2019). Despite its associated plant damage, *N. tenuis* is widely used across South-eastern Spain as it is an efficient agent against the various pests it controls (Arnó et al., 2010). Furthermore, the aforementioned phytophagy has been demonstrated to have benefits by triggering predator-induced defences, including attracting parasitoids, repulsing other herbivorous pests, and restricting accumulation of viruses (Bouagga et al., 2019; Pérez-Hedo et al., 2018, 2015).

In recent years, the controversial success of *N. tenuis* has encouraged the scientific community to study this predatory mirid (Puentes et al., 2018). However, some issues remain to be addressed, such as the genetic variation in commercial stocks of similar biological control agents when compared to wild populations, with the former often diminished in comparison to the latter as seen in other biological control agents (Paspati et al., 2019; Rasmussen et al., 2018; Streito et al., 2017). In order to compare biological control stock to wild (or wild-caught) populations, determining the current diversity and genetic variation of the commercial stock is important. Finally, *N. tenuis* is known to host bacterial symbionts, including *Wolbachia* and *Rickettsia*, though the effect of these bacteria on their host is relatively unknown (Caspi-Fluger et al., 2014). Sequence data can provide additional insight into potential symbionts as well as identify potential LGTs (lateral gene transfers) between host and symbiont.

With all of these fascinating avenues of research in mind, it may be surprising to learn that, aside from a mitogenome (Dai et al., 2012), a regional population analysis (Xun et al., 2016), and more recent work shedding light on evidence of LGT (P. Xu et al., 2019), little genomic information exists for *N. tenuis* and there is no published *N. tenuis* genome. Characteristics such as karyotype, sex chromosome system, and presence or absence of telomeric repeats are currently unknown. A likely reason for this absence of genomic resources is that advances made in sequencing technology are often juxtaposed to the complexities of insect life cycles and difficulties in obtaining enough high quality genomic material due to size and exoskeleton (Leung et al., 2019; Richards and Murali, 2015). Additionally, current assembly tools have a hard time dealing with heterozygosity; therefore, a genome assembly is benefited by sequencing material of reduced genetic heterozygosity for a more contiguous assembly. Reduced heterozygosity is often difficult to achieve in diploid insects where the genetic variation within a population is unknown or the species cannot be inbred (Keeling et al., 2013).

Generating the genomes of highly heterozygous, diploid, and relatively small insects is tricky; researchers have to be prepared to balance their expectations and the available technology (Ellegren, 2014; Leung et al., 2019). While a single diploid individual may yield enough material for an Illumina-only library, assembly may be difficult due to large repeat regions that extend beyond the insert size of the library. Conversely, enough material could be obtained for sequencing on a long-read platform, but may require pooling material from multiple individuals, potentially complicating assembly due to the heterozygosity of the population. While possible solutions include estimating the heterozygosity or setting up inbred populations (which can be nearly impossible if deleterious effects of inbreeding need to be avoided or if the presence of a complementary sex determining system limits inbreeding (Szűcs et al., 2019; van Wilgenburg et al., 2006)), an alternative is to create a linked-read library. The 10x Genomics platform creates a microfluidic partitioned library that individually barcodes minute amounts of long strands of DNA for further amplification (10x Genomics Inc., Pleasanton, CA, USA). This library is then sequenced on a short-read sequencing platform and then assembled using the barcodes to link reads together into the larger fragment (i.e. Chin et al. 2016; Jones et al. 2017). This method allows for a library to be constructed from a single individual that contains additional structural information to aid assembly (such as phasing), removing the need for pooling multiple individuals and avoiding assembly difficulties in repetitive regions. Additional information, such as karyotype, can further improve genomes in the assembly stage as well as inform further directions of research by providing chromosome-level context, encouraging further improvement of a genome beyond its initial release.

Here we present the genome of *Nesidiocoris tenuis* achieved by sequencing a linked-read library of a single adult female bug, along with an annotation based on transcriptome, homology-based, and *ab initio* predictions. In addition to the genome, various avenues for future research are initiated to raise the profile of *N. tenuis* as a research organism, including cytogenetic analyses, protein cluster analysis, and a genome-wide pooled sequencing population genetics analysis. These resources benefit biological control research, as more knowledge becomes available to use in research as well as knowledge of the species for taxonomic and phylogenetic purposes.

## Methods

### Species origin and description

Individuals of *N. tenuis* were received either from the commercial biological control stock at Koppert Biological Systems, S. L. (Águilas, Murcia, Spain) (KBS) or from the population maintained for less than a year at Wageningen University and Research (WUR) Greenhouse Horticulture (Bleiswijk, The Netherlands), which in turn were originally sourced from the KBS commercial population. Material used for DNA sequencing, PCR testing, pooled sequencing, and cytogenetics was from the KBS population, while material used for RNA sequencing was from the WUR Greenhouse Horticulture population. Additional species used for cytogenetic comparison purposes were sourced from two separate laboratory populations within the Biology Centre CAS in České Budějovice, Czech Republic: *Triatoma infestans* (Klug) (Hemiptera: Reduviidae) individuals were obtained from a laboratory colony at the Institute of Parasitology that was originally sourced from Bolivia (Schwarz et al., 2014), while *Ephestia kuehniella* (Zeller) (Lepidoptera: Pyralidae) individuals were obtained from a wild-type laboratory colony at the Institute of Entomology (Marec and Shvedov, 1990). Species identification of the KBS population was confirmed via COI sequencing using a PCR amplification protocol (Itou et al., 2013), in addition to testing for the presence of *Wolbachia* via PCR amplification protocol (Zhou et al., 1998).

### Flow cytometry

Genome size was estimated with flow cytometry on propidium-iodide stained nuclei. Individuals from a mixed *Drosophila melanogaster* (Meigen) (Diptera: Drosophilidae) laboratory population (May et al., 2019) were used as the standard for genome size comparison. Following established preparation protocols (De Boer et al., 2007), three samples of single *D. melanogaster* heads, two samples of single *N. tenuis* heads, and one sample of a single *N. tenuis* head pooled with a single *D. melanogaster* head were analysed in a FACS flow cytometer (BD FACSAria^TM^ III Fusion Cell Sorter, BD Biosciences, San Jose, USA). With the known genome size of *D. melanogaster* of 175 Mbp, we could calculate an approximate genome size relative to the amount of fluorescence (Hare and Johnston, 2011).

### gDNA Extraction

A single female *N. tenuis* was placed in a 1.5 mL safelock tube with 5-8 one mm glass beads and frozen in liquid nitrogen and shaken for 30 s in a Silamat S6 shaker (Ivoclar Vivadent, Schaan, Liechtenstein). DNA was then extracted using the Qiagen MagAttract Kit (Qiagen, Hilden, Germany). Following an overnight lysis step with Buffer ATL and proteinase K at 56°C, extraction was performed according to MagAttract Kit protocol. Elution was performed in two steps with 50 µL of Buffer AE (Tris-EDTA) each time, yielding 424 ng of genomic DNA (gDNA) in 100 µL as measured with an Invitrogen Qubit 2.0 fluorometer using the dsDNA HS Assay Kit (Thermo Fisher Scientific, Waltham, USA).

### Library Preparation and Sequencing

Following extraction, gDNA was further diluted to 1 ng/µl following the Chromium Genome Reagent Kits Version 1 User Guide (version CG-00022) (10x Genomics, Pleasanton, USA). A library of Genome Gel Beads was combined with 1 ng of gDNA, Master Mix, and partitioning oil to create Gel Bead-In-EMulsions (GEMs). The GEMs underwent an isothermal amplification step and barcoded DNA fragments were recovered for Illumina library construction (Illumina, San Diego, USA). The library was then sequenced on an Illumina HiSeq 2500 at the Bioscience Omics Facility at Wageningen University and Research (Wageningen, The Netherlands), yielding 212,910,509 paired-end reads with a read length of 150 bp. The first 23 bp of each forward read is a 10X GEM barcode used in the assembly process. Forward read quality was similar to that of the reverse reads, and no reads were flagged for poor quality in a FastQC assessment (Andrews et al., 2015).

### Assembly

Using the reads, a k-mer count analysis was performed using GenomeScope on k-mer sizes of 21 and 48, which was used to infer heterozygosity (Vurture et al., 2017). Assembly was performed using all available reads with the GEM barcodes incorporated during the Chromium library preparation in Supernova v2.1.1 (10X Genomics, Pleasanton, USA), with default settings (Weisenfeld et al., 2017). This assembly, v1.0, underwent a preliminary decontamination using NCBI BLASTn v2.2.31+ against the NCBI nucleotide collection (nt) focusing on scaffolds with over 95% homology to bacteria (Camacho et al., 2009), followed by the more elaborate method described below (Detecting contamination and LGT events). Finally, 100% duplicate scaffolds were identified using the *dedupe* tool within BBTools (sourceforge.net/projects/bbmap/), and removed alongside the contaminated scaffolds, resulting in assembly v1.5. Attempts at further deduplication by adjusting the threshold (such as 95% duplication) resulted in further deletions, but at larger scaffold size, percentage is a rather blunt tool and any percentage is an arbitrary cut-off, so we decided to only remove true duplicates. Assembly completeness for both assemblies were determined using BUSCO v3.0.2 and the insect_odb9 ortholog set (Simão et al., 2015), while assembly statistics were determined using QUAST (Gurevich et al., 2013).

### Detecting contamination and LGT events

Lateral gene transfers (LGTs) from bacteria into metazoan genomes were once thought to be rare or non-existent, but are now known to be relatively common and can evolve into functional genes (Dunning Hotopp et al., 2007; Husnik and McCutcheon, 2018). We therefore screened our insect genome for LGTs from bacteria. As insect genome assemblies often contain scaffolds from associated bacteria, we first screened for such “contaminating” scaffolds and moved them into a separate metagenomic multiFASTA (S1.3.4).

We used a DNA-based computational pipeline to both identify likely contaminating bacterial scaffolds in the assembly, and to detect potential LGT from bacteria into the insect genome. The LGT Pipeline was modified from an earlier version developed by David Wheeler and John Werren (Wheeler et al., 2013), and has been used to screen for bacterial “contamination” and LGTs in a number of arthropod genomes before (e.g. bedbug *Cimex lectularis* L. (Hemiptera: Cimicidae) (Benoit et al., 2016), parasitoid wasp *Trichogramma pretiosum* Riley (Hymenoptera: Trichogrammatidae) (Lindsey et al., 2018), and the milkweed bug *Oncopeltus fasciatus* (Dallas) (Hemiptera: Lygaeidae) (Panfilio et al., 2019)). In some cases, entire or nearly complete bacterial genomes have been retrieved from arthropod genome projects (e.g. (Benoit et al., 2016; Lindsey et al., 2016)).

#### Detection of Bacterial Scaffolds in the Assembly

To detect bacterial contaminating scaffolds, the following method was used after the preliminary bacterial contamination assessment described above. First, each scaffold was broken into 1 kbp fragments and each fragment was subsequently searched with BLASTn against an in-house reference database that contains 2,100 different bacterial species (complete list in S1.3.1) which was masked for low complexity regions using the NCBI Dustmasker function (Morgulis et al., 2006). We recorded each bacterial match with bitscore > 50, the number of bacterial matches, total bacterial coverage in the scaffold, proportion of the scaffold covered, total hit width of coverage (the distance between the leftmost and rightmost bacteria hit proportional to the scaffold size) and the bacterial species with the greatest number of matches within the scaffold from the in-house bacterial data base. It should be noted that the latter method does not indicate the actual bacterial species from which the scaffold was derived, as it is based on similarity to a curated database – that determination would require follow-up analysis, which was not performed in this study.

Any criterion for deciding whether a scaffold comes from a bacterium is unavoidably arbitrary: Too stringent and insect scaffolds are included; too lax and insect scaffolds are inappropriately removed. We applied a cut-off of ≥ 0.40 proportion bacterial hit width, which has performed well to remove contamination in a few test cases where we have manually examined scaffolds near the cut-off. All instances of contaminated scaffolds were removed from the assembly and are available in supplementary materials as a list (S1.3.2) and a multi-FASTA file (S1.3.3).

#### Identifying LGT Candidate Regions

We used the same DNA based computational pipeline to identify potential LGTs from bacteria into the insect genome. The basic method is as follows: as before, scaffolds from the genome assembly are broken into 1 kbp intervals, which are searched against a bacterial genome database. Any positive bacterial hit in a 1 kbp region (bitscore > 50) was then searched against a database containing transcripts from the following eukaryotes: *Xenopus*, *Daphnia*, *Strongylocentrotus*, *Mus*, *Homo sapiens*, *Aplysia*, *Caenorhabditis*, *Hydra*, *Monosiga,* and *Acanthamoeba* (ftp://ftp.hgsc.bcm.edu/I5K-pilot/LGT_analysis/All_species_genomes/lgt_finder_blastn_database_directories/). The purpose of this eukaryotic screening is to identify highly conserved genes that are shared between eukaryotes and bacteria and exclude these from further analysis. To focus our attentions on the most likely LGT candidates, we selected hits with a bitscore = 0 in the corresponding reference eukaryote database and bitscore > 75 from the bacterial database. We also screened the output for adjacent 1 kbp pieces that contain bacterial matches and reference eukaryote bitscore = 0 and fused these adjoining pieces for analysis.

LGT candidate regions were then manually curated as follows: each candidate region was searched with BLASTn to the NCBI nr/nt database. If this search indicated that the region’s nucleotide sequence was similar or identical to the nucleotide sequence of a known gene in related insects, it was discarded as a likely conserved insect gene. Regions were retained only when the matches to other insects were sporadic, as our experience has indicated that these can be independent LGTs into different lineages. If no match was found, the region was additionally searched with BLASTx to the NCBI nr/nt database. If this second search also resulted in no hits to multiple insect proteins, it was called an LGT candidate. In this case, we additionally identified the best bacterial match using the NCBI nr and protein databases. Using the gene annotation information, we then evaluated the flanking genes within the scaffold to determine whether they were eukaryotic or bacterial, we determined whether the LGT region was associated with an annotated gene within the insect genome, and we observed with transcriptome data if RNA sequencing data showed evidence of transcriptional activity in the LGT region. This short list is available in the supplementary materials (S1.3.4.)

### RNA extraction, library construction, and sequencing

Juveniles, adult males, and adult females (approximately 4-5 of each) were prepared for RNAseq using the RNeasy Blood and Tissue Kit (Qiagen). Individuals were placed in a 1.5 mL safelock tube along with 5-8 one mm glass beads placed in liquid nitrogen and then shaken for 30 s in a Silamat S6 shaker (Ivoclar Vivadent). RNeasy Blood and Tissue Kit (Qiagen) was used according to manufacturer’s instructions. Samples were assessed for quality and RNA quantity using an Invitrogen Qubit 2.0 fluorometer and the RNA BR Assay Kit (Thermo Fisher Scientific). These three RNA samples were then processed by Novogene Bioinformatics Technology Co., Ltd., (Beijing, China) using poly(A) selection followed by cDNA synthesis with random hexamers and library construction with an insert size of 550-600 bp. Paired-end sequencing was performed on an Illumina HiSeq 4000 according to manufacturer’s instruction.

### Gene finding, transcriptome assembly, and annotation

For the *ab initio* gene finding, a training set was established using the reference genome of *D. melanogaster* (Genbank: GCA_000001215.4; Release 6 plus ISO1 MT) and the associated annotation. The training parameters were used by GlimmerHMM v3.0.1 for gene finding in the *N. tenuis* genome assembly v1.5 (Majoros et al., 2004). For homology-based gene prediction, GeMoMa v1.6 was used with the *D. melanogaster* reference genome alongside our RNAseq data as evidence for splice site prediction (Keilwagen et al., 2016). For evidence-based gene finding, each set of RNAseq data (male, female, and juvenile) was mapped to the *N. tenuis* genome separately with TopHat v2.0.14 with default settings (Trapnell et al., 2009). After mapping, Cufflinks v2.2.1 was used to assemble transcripts (Trapnell et al., 2010). CodingQuarry v1.2 was used for gene finding in the genome using the assembled transcripts, with the strandness setting set to ‘unstranded’ (Testa et al., 2015).

The tool EVidenceModeler (EVM) v1.1.1 was used to combine the *ab initio*, homology-based, and evidence-based information, with evidence-based weighted 1, *ab initio* weighted 2, and homology-based weighted 3 (Haas et al., 2008). The resulting amino acid sequences were searched with BLASTp v2.2.31+ on a custom database containing all SwissProt and Refseq genes of *D. melanogaster* (Acland et al., 2014; Boutet et al., 2008; Camacho et al., 2009). The top hit for each amino acid sequence/gene was retained and its Genbank accession number and name are found within the annotation. If no hit was found, an additional search in the NCBI non-redundant protein database (nr) was performed to obtain additional homology data.

### Functional annotation and GO term analysis

Gene attributes from the annotation were used to construct a list of genes to be used in Gene Ontology (GO) term classification. Duplicate accession numbers were removed alongside cases where no BLAST hit was found. The remaining accession IDs were converted into UniProtKB accession IDs using the UniProt ID mapping feature (Huang et al., 2011). These UniProtKB accession IDs were in turn used with the DAVID 6.8 Functional Annotation Tool to assign GO terms to each accession ID with the *D. melanogaster* background and generate initial functional analyses (Huang et al., 2009a, 2009b).

### Ortholog cluster analysis and comparison

The complete gene set of *Nesidiocoris tenuis* was compared to those of three additional hemipteran species: the bed bug *Cimex lectularis*, the brown marmorated stinkbug *Halyomorpha halys* (Stål) (Hemiptera: Pentatomidae), and the pea aphid *Acyrthosiphon pisum* (Harris) (Hemiptera: Aphididae) using OrthoVenn2 (L. Xu et al., 2019). The gene set of *A. pisum* is the 2015 version from AphidBase (Legeai et al., 2010; Richards et al., 2010) as maintained on the OrthoVenn2 server. The *H. halys* 2.0 complete gene set was used (Lee et al., 2009) along with the complete gene set of *C. lectularis* (Clec 2.1, OGSv1.3) (Benoit et al., 2016; Thomas et al., 2020), both of which were retrieved from the i5K Workspace (Poelchau et al., 2015). An ortholog cluster analysis was performed on all four gene sets via OrthoVenn2 with the default settings of E-values of 1e-5 and an inflation value of 1.5.

### Cytogenetic analysis

#### Slide preparations

To determine karyotype, *N. tenuis* individuals were obtained from the KBS population and prepared for cytogenetic experiments. Chromosomal preparations were prepared from the female and male reproductive organs of adults and juveniles by spreading technique according to Traut (1976) with modifications from Mediouni *et al*. 2004 (Mediouni et al., 2004; Traut, 1976). After inspection via stereomicroscope to confirm presence of chromosomes, slides were dehydrated in ethanol series (70, 80, and 100%, 30 s each) and stored at −20°C for future use.

#### 18S rDNA probe preparation and Fluorescence In Situ Hybridisation (FISH)

To confirm the presence of 18S rDNA sequences in the assembled genome, the previously published partial 18S rDNA sequence of *N. tenuis* (GU194646, Jung and Lee, 2012) was used as a BLAST query against the *N. tenuis* v1.5 genome. To verify sequence homology, the obtained 18S rDNA sequences were subsequently compared to the previously published sequence.

For preparation of the probe, we isolated gDNA from two *N. tenuis* females with the NucleoSpin DNA Insect Kit (Macherey-Nagel, Düren, Germany) according the manufacturer’s protocol. gDNA was used as template in PCR to amplify the 18S rDNA sequence using primers 18S-1 and 18S-4 as described in Jung and Lee (2012). Obtained products were purified using the Wizard SV Gel and PCR Clean-Up System (Promega, Madison, WI, USA), and subsequently cloned using the pGEM-T Easy Vector System (Promega, Madison, WI) according to the manufacturer’s protocol. Plasmids were extracted from positive clones with the NucleoSpin Plasmid kit (Macherey-Nagel) following the manufacturer’s protocol, confirmed by sequencing (SEQme, Dobříš, Czech Republic), and used as template in PCR with the 18S-1 and 18S-4 primers. PCR-products were purified, and used as template for labelling by a modified nick translation protocol as described by Kato *et al*. (2006) with modifications described in Dalíková *et al*. (2017), using biotin-16-dUTP (Jena Bioscience, Jena, Germany) and an incubation time of 35 minutes at 15°C (Dalíková et al., 2017a; Kato et al., 2006). Fluorescence *in situ* hybridization (FISH) was performed as described in Sahara *et al*. (1999) with modifications described in Zrzavá *et al*. (2018) (Sahara et al., 1999; Zrzavá et al., 2018).

#### Sex chromosome identification

Determination of the sex chromosome constitution is important for the assembly of the *N. tenuis* genome to identify any potential missing information due to sequencing a single sex, as well as add to knowledge on sex chromosomes in Miridae. Comparative Genomic Hybridization (CGH), and Genomic *In Situ* Hybridization (GISH) were, therefore, used to identify the sex chromosomes of *N. tenuis*. The reproductive organs of adult females were dissected out to avoid potential male gDNA contamination, as the mated status was unknown, after which remaining tissue was snap-frozen in liquid nitrogen and stored at −20°C until further use. Adult males were not dissected but otherwise treated the same. Female and male gDNA was extracted from 10-20 pooled individuals using cetyltrimethylammonium bromide (CTAB) gDNA isolation with modifications (Doyle and Doyle, 1990). Samples were mechanically disrupted in extraction buffer (2% CTAB, 100 mM Tris-HCl pH 8.0, 40 mM EDTA, 1.4 M NaCl, 0.2% β-mercaptoethanol, 0.1 mg/mL proteinase K), and incubated overnight at 60°C with light agitation. An equal volume chloroform was added, tubes were inverted for 2 min, and samples were centrifuged 10 min at maximum speed. The aqueous phase was transferred to a new tube, RNase A (200 ng/µL) was added and samples were incubated 30 min at 37°C to remove RNA. DNA was precipitated by adding 2/3 volume isopropanol, gently inverting the tubes, and centrifugation for 15 min at maximum speed. Pellets were washed twice with 70% ethanol, air-dried briefly, and dissolved overnight in sterile water. DNA was stored at −20°C until further use. Probes were prepared with 1 μg gDNA using Cy3-dUTP (for female gDNA), or fluorescein-dUTP (for male gDNA) (both Jena Bioscience, Jena, Germany) by nick translation mentioned above with an incubation time of 2-2.5 hours at 15°C. CGH and GISH were performed according to Traut et al. (1999) with modifications described in Dalíková et al. (2017) (Dalíková et al., 2017b; Traut et al., 1999).

#### Detecting a telomeric motif

Initially, we searched both the raw sequencing data and the assembled genome for presence of the ancestral insect telomere motif (TTAGG)*_n_*, which is known to be absent in several Miridae (Grozeva et al., 2019; Kuznetsova et al., 2011), and tested for its presence using Southern dot blot in *N. tenuis*. gDNA was isolated from *N. tenuis*, and positive controls *E. kuehniella* and *T. infestans*, using CTAB DNA isolation described above. DNA concentrations were measured by Qubit 2.0 (Broad Spectrum DNA Kit) (Invitrogen) and diluted to equalize concentrations, after which 500 ng and 150 ng of each specimen was spotted on a membrane and hybridized as described in (Dalíková et al., 2017b). As a negative control, an equal amount of sonicated DNA from the chum salmon *Oncorhynchus keta* (Walbaum) (Salmoniformes: Salmonidae) (Sigma-Aldrich, St. Louis, MO, USA), was spotted on the same membrane. Probe template was prepared using non-template PCR according to Sahara *et al*. (1999) and labelling with digoxigenin-11-dUTP (Jena Bioscience) was performed using nick translation, with an incubation time of 50 min according to Dalíková *et al*. (2017) (Dalíková et al., 2017a; Sahara et al., 1999). Absence of the insect telomere motif (TTAGG)*_n_* was confirmed by dot blot, and three sequence motifs, (TATGG)*_n_*, (TTGGG)*_n_*, and (TCAGG)*_n_*, were selected as potential telomeric motifs in *N. tenuis* based on high copy numbers in the genome and sequence similarity to the ancestral insect telomere motif. Copy numbers were determined by Tandem Repeat Finder (TRF, version 4), on collapsed quality filtered reads corresponding to 0.5x coverage with default numeric parameters except maximal period size, which was set to 25 bp (Benson, 1999). TRF output was further analysed using Tandem Repeat Analysis Program (Sobreira et al., 2006). Probe template, and subsequent labelling of the probes, was done as described above with slight alterations. To obtain optimal length of fragments for labelling, non-template PCR was performed with reduced primer concentrations (50 nM for each primer). In addition, probes were labelled by biotin-16-dUTP (Jena Bioscience) using nick translation as described above, with an incubation time of 50 min. FISH was performed as described above for 18S rDNA.

#### Repeat identification and visualization

To assess the repetitive component of the *N. tenuis* genome, we used RepeatExplorer, version 2, on trimmed and quality-filtered reads with default parameters (Novák et al., 2013). Repeats with high abundance in the genome were selected and amplified using PCR. These products, named Nt_rep1, were additionally cloned and template for probe labelling was prepared from plasmid DNA as described above, see 18S rDNA probe preparation. Probes were labelled by PCR in a volume of 25 µL consisting of 0.625 U Ex *Taq* polymerase (TaKaRa, Otsu, Japan), 1x Ex *Taq* buffer, 40 µM dATP, dCTP, and dGTP, 14.4 µM dTTP, 25.6 µM biotin-16-dUTP (Jena Bioscience), 400 nM of forward and reverse primer, and 1 ng of purified PCR-product. The amplification program consisted of an initial denaturing step at 94°C for 3 min, followed by 35 cycles of 94°C for 30 s, 55°C for 30 s, 72°C for 1 min, and a final extension at 72°C for 2 min. The FISH procedure was performed as described for 18S rDNA. Abundance and distribution of Nt_rep1 in the assembled genome was assessed using NCBI Genome Workbench version 2.13.0. The complete list of primers used in this study can be found in Table S1.6.2.

### Pooled sequencing and population analysis

For this project, it was important to use existing data wherever possible to test the utility of the genome and possible research avenues. Therefore, we analysed whole genome sequence data originally generated for another *N. tenuis* genome assembly project that has not been published before. In the original set-up, ten females were collected from the KBS population for pooled sequence analysis. DNA was isolated from this pooled cohort using the “salting-out” method as described in Sunnucks and Hales with a final volume of 20 µL (Sunnucks and Hales, 1996) and then treated with 2 µL RNase. The paired-end library was sequenced on an Illumina HiSeq2500 platform by Macrogen Inc. (Seoul, Korea) with read sizes of 100 bp. Reads were assessed for quality using FastQC (Andrews et al., 2015) and adapters were trimmed with Trimmomatic (Bolger et al., 2014). Following quality filtering, reads with phred scores lower than 20 were discarded. Heterozygosity was calculated using jellyfish v2.3.0 and GenomeScope v1.0 with a k-mer size of 21 and default parameters (Marçais and Kingsford, 2011; Vurture et al., 2017).

Instead of genome assembly, these whole genome sequence reads were adjusted and subsequently used in a pooled sequencing (pool-seq) population analysis with our genome. First, the reads were randomly subsampled to a coverage of 10X (in a pool with 10 females, this results in approximately 1X coverage per female) using CLC Genomics Workbench 12 (Qiagen). Using the PoPoolation v1.2.2 pipeline (Kofler et al., 2011), these reads were aligned to an adapted v1.5 genome, where scaffolds smaller than 10,000 bp were removed, and aligned reads were binned into windows using the bwa and samtools packages (Li et al., 2009). Pileup files and the scripts from the PoPoolation pipeline were used to produce variance sliding windows analyses of neutrality, *Tajima’s D*, and nucleotide diversity, *Tajima’s pi* (π) with default settings and a pool size of 40. Window and step sizes of both 10,000 and 5,000 were tested, as well as using the “basic-pipeline/mask-sam-indelregions.pl” pipeline to mask indel regions of the SAM file – this ensures that indel regions are not calculated. Of the 18,000,000 reads, 817,226 had regions of indels masked.

## Results

### Species origin, description, and data availability

The presence of *Wolbachia* in the KBS biological control stock was confirmed (Electrophoresis gel image in supplementary material, S1.1). All sequence data generated, including raw reads, assembly, and annotation, can be found in the EMBL-EBI European Nucleotide Archive (ENA) under BioProject PRJEB35378. An additional, complete annotation file (.gff) is also available (Ferguson, 2020).

### Genome assembly and size

The single adult female *N. tenuis* yielded 424 ng total DNA. The 10X Genomics Chromium reaction and subsequent Illumina sequencing resulted in more than 212 million paired-end reads. The inferred heterozygosity, based on GenomeScope, was between 1.675% and 1.680% for a k-mer size of 21, and between 1.250% and 1.253% for a k-mer size of 48. Genome size estimates at this point were between 306 Mbp (k-mer=21) and 320 Mbp (k-mer=48). Following assembly with Supernova, assembly v1.0 was approximately 388 Mbp in size and comprised of 44,273 scaffolds (5.91% ambiguous nucleotides).

Assembly v1.0 was then assessed for contamination with a preliminary search against the NCBI for bacterial homology (see below for more details). Several scaffolds with high amounts of bacterial sequence contamination were identified, indicating that further decontamination of the assembly was required. A decontamination pipeline was used to identify and remove a total of 3,043 scaffolds, while those identified as potential examples of LGT were kept. From the remainder, an additional 4,717 were identified as being identical duplicates and were removed. At this point, the resulting assembly was finalised and designated v1.5. This assembly is 355 Mbp in size, consisting of 36,513 scaffolds (6.29% ambiguous nucleotides). Quality and completeness of v1.5 using BUSCO indicated a completeness of 87.5% (65.6% single copy orthologs, 21.9% duplicated orthologs), while 7.1% orthologs were fragmented and 5.4% were missing (n=1658).

Initially, the genome size of *N. tenuis* was estimated by flow cytometry to be 232 Mbp, with a confidence interval of 20 Mbp (See supplementary material S1.2 for more details). Further estimates via k-mer analysis of sequence data in GenomeScope indicated an expected genome size of 306 Mbp (k-mer=21) or 320 Mbp (k-mer=48). Both the flow cytometry and sequence data estimates are smaller than the 355 Mbp of the final assembly (v1.5). In total, the *N. tenuis* genome has 36,513 scaffolds, with the largest scaffold being 1.39 Mbp, though the majority of scaffolds are under 50,000 bp in size. The number of gaps per 100 kbp is 6292.10 (6.29% of the genome). Details on the assemblies can be found in Table 1.

**Table 1.** Assembly statistics for both versions of the *Nesidiocoris tenuis* assembly, pre- and post-decontamination

| Assembly Version | Size (bp) | No. of Scaffolds | N50 (bp) | Largest scaffold (bp) | No. of N's per 100 kbp (% of genome) | BUSCO score, Complete% (Single%, Duplicate%) |
| --- | --- | --- | --- | --- | --- | --- |
| 1.0 | 387,724,797 | 44,273 | 27,195 | 1,392,896 | 5912.60 (5.91) | 81.3 (60.6, 20.7) |
| 1.5<br>(final assembly) | 355,120,802 | 36,513 | 28,732 | 1,392,896 | 6292.10 (6.29) | 87.5 (65.6, 21.9) |

### Assessment of potential symbionts and LGT candidates

#### Potential symbionts

The initial assembly (v1.0) was decontaminated using two bacterial decontamination pipelines: the first pipeline broadly utilised BLASTn to identify scaffolds with high amounts of bacterial sequences against the NCBI nr database, while the second pipeline is more specified and uses BLASTn against a list of known contaminants and symbionts (S1.3) and is adapted from previous work (Wheeler et al., 2013). The first decontamination pipeline identified and removed 1,443 scaffolds with high bacterial content, and the second decontamination pipeline identified and removed an additional 1,600 scaffolds alongside potential LGT events. All removed scaffolds are available in S1.3. The hits from the second pipeline were used to create a list of potential contaminants or symbionts of this particular *N. tenuis* individual used for whole genome sequencing according to genus, base pair content, and number of scaffolds affected (Table 2). The majority of these scaffolds (1470) are under 5 kbp in length, with an additional 61 scaffolds falling between 5-10 kbp. The ten largest scaffolds are putatively associated with *Pantoea* and relatives (three of 561,7472 bp, 205,621 bp, and 131,905 bp), *Sodalis* (326,101 bp), *Erwinia* (254,660 bp; 220,307 bp; 154,581 bp), and *Citrobacter* (239,269 bp; 190,839 bp). We emphasize that these “calls” are very preliminary, as they are based on the most frequent hits in the bacterial matches in each scaffold, rather than comprehensive gene annotations. Nevertheless, they do indicate a range of bacterial types associated with *N. tenuis*, and the scaffold assemblies are likely to contain some complete or near complete bacterial genomes of interest.

**Table 2.** Genera of potential symbionts or contaminants as determined by decontamination pipeline based on known contaminants and symbionts against *Nesidiocoris tenuis* assembly v1.0. Identification is according to largest hit percentage, multiple bacterial sections possible in each scaffold. Affected scaffolds were removed leading to assembly v1.5 and are available in S1.2. Full list of hits available in supplementary material S1.4.

| **Bacteria Genus** | **Total amount in genome (bp)** | **Number of scaffolds affected** |
| --- | --- | --- |
| *Pantoea* | 1,379,962 | 307 |
| *Erwinia* | 1,342,298 | 456 |
| *Citrobacter* | 349,562 | 40 |
| *Rickettsia* | 204,934 | 50 |
| *Sodalis* | 189,471 | 17 |
| *Cronobacter* | 180,427 | 33 |
| *Wolbachia* | 137,109 | 47 |
| *Enterobacter* | 110,104 | 70 |
| *Serratia* | 100,197 | 49 |
| *Klebsiella* | 44,248 | 30 |
| *Pseudoalteromonas* | 39,779 | 118 |
| *Pectobacterium* | 28,130 | 16 |
| *Shigella* | 25,221 | 24 |
| *Yersinia* | 24,052 | 21 |
| *Dickeya* | 23,312 | 18 |
| *Salmonella* | 19,243 | 15 |
| *Photorhabdus* | 18,283 | 14 |
| *Escherichia* | 16,301 | 11 |
| *Rahnella* | 12,479 | 10 |
| *Burkholderia* | 7,825 | 17 |
| *Xenorhabdus* | 6,406 | 8 |
| *Arsenophonus* | 5,070 | 8 |
| *Pseudomonas* | 3,427 | 2 |
| *Vulcanisaeta* | 3,420 | 3 |
| *Ralstonia* | 3,370 | 7 |
| *Paenibacillus* | 3,070 | 8 |
| Others (105) | 56,287 | 201 |
| **Total** | **4,333,987** | **1,600** |

Sorting scaffolds across the range of bacterial genera matches gives 131 genera with some substantial representation: *Erwinia* (2,078,531 bp), *Pantoea* (2,226,778 bp), *Citrobacter* (594,902 bp), *Sodalis* (355,847 bp), *Cronobacter* (314,511 bp), and *Rickettsia* (483,217 bp) (Table 2). In addition to known symbiont *Rickettsia*, previously established via PCR and known symbiont *Wolbachia* is also present in the results (137,109 bp) (Table 2). Multiple genera of bacteria can be found on a single scaffold, likely due to misassembly. The full list of bacterial scaffolds and multiFASTA file is available with details in supplementary materials S1.3.2 and S1.3.3.

#### LGT Candidates

We continued with our detection of potential LGT events by further assessing a handful of strong candidates. Two of these regions occur on scaffolds 22012 and 22013, which are of similar length (22,634 bp and 22,957 bp, respectively) and are highly similar on a nucleotide level. Scaffold 22013 appears to have additional nucleotides on each flanking side, with some indels and SNPs between the two scaffolds. The putative LGT region in question is belonging to or gained from a *Sodalis* species, coding for phenazine biosynthesis protein *PhzF* (OIV46256.1). This region also showed transcriptional support, and is flanked by conserved insect genes, most immediately *Rab19* (NP_523970.1) on one side and an uncharacterized protein, Dmel_CG32112 (NP_729820.2), on the other side. Two additional LGTs were found in the current assembly that match *Rickettsia* sequences (scaffolds 4712 and 27281), which contain a segment of the rickettsial genes *elongation factor G* and *AAA family ATPase* genes, respectively. One corresponds to a gene model, while the other does not, and there is no evidence of expression for either in the current male, female, and mixed sex juvenile RNA sequencing data. More information on these candidate regions can be found in S1.3.4.

### *Ab initio* gene finding, transcriptome assembly, and annotation

To obtain a comprehensive set of transcripts for *N. tenuis*, three separate libraries of multiple individuals were prepared – males, females, and juveniles from different stages of mixed sex. More than 77 million 150 bp pair-end reads were generated. Filtering the reads for quality led to a slightly reduced total of 76,711,096 paired reads (male: 28,413,231 paired reads; female: 24,075,901 paired reads; juvenile: 24,221,964 paired reads) to be used for evidence-based gene finding. The mapping and assembling of reads of the three individual samples as well as the pooled reads resulted in four transcriptomes: male, female, juvenile, and the combined transcriptome.

The male, female, juvenile, and combined annotations from the evidence-based gene finding was used alongside homology-based findings and *ab initio* annotations in a weighted model, resulting in complete annotations for the assembly. When gene name assignment via the SwissProt database resulted in “no hit,” tracks are named “No_blast_hit.” This occurred in 1,556 mRNA tracks and represents approximately 6% of the official gene set. The majority of tracks were annotated with reference to SwissProt or GenBank accession number of the top BLASTp hit.

CodingQuarry predicted 56,309 genes from the mapped transcript evidence, while *ab initio* gene finding using GlimmerHMM resulted in 39,888 genes and homology-based gene finding with GeMoMa resulted in 6,028 genes. The complete gene set for *N. tenuis* was created using EVidenceModeler, where a weighted model using all three inputs resulted in a complete gene set of 24,688 genes.

### Functional annotation and GO term analysis

The complete gene set of 24,668 genes was deduplicated and genes with no correlating BLASTp hit were removed. The remaining 11,724 genes were mapped to UniProtKB IDs, resulting in 11,261 genes with a matching ID after another round of deduplication (80 duplicates found). The remaining 383 genes either did not match to a UniProt KB ID or were considered obsolete proteins within the UniParc database. DAVID used 8,920 genes for the functional annotation analysis, of which 78.4% (6503) contribute to 19 biological processes, 75.8% (6826) contribute to 100 different cellular components, and 72.8% (6032) contribute to 91 categories of molecular functions (genes can code to multiple GO terms). The remaining genes were uncategorized. Data linking the genes to the GO terms, the DAVID Gene List Report, and the DAVID Gene Report are available in S1.4.

### Ortholog cluster analysis

The complete gene set of *N. tenuis* was compared to those of three additional species: the bed bug *C. lectularis*, the brown marmorated stinkbug *H. halys*, and the pea aphid *A. pisum* using OrthoVenn2. The ortholog analysis summary is presented in Table 3 and visualized in Figure 1. *N. tenuis* has a similar number of clusters (8,174) as compared to *C. lectularis, H. halys and A. pisum* (7,989; 9,584; and 8,765, respectively). In total 14,512 clusters are assigned, 12,964 of which are orthologous clusters (contains at least two species), and the remaining 1,548 are single-copy gene clusters. There are 9,136 singleton clusters in *N*. *tenuis*, 3,573 in *C. lectularis*, 2,170 in *H. halys*, and 7,298 in *A. pisum*. The amount of singleton clusters, i.e. proteins that do not cluster, indicate that *N. tenuis* differs the most from the other species, as 37.04% of the proteins are singletons. Just over half of the orthologs cluster with *N. tenuis*, where 6,338 clusters are outside of *N. tenuis* as compared to the 8,174 clusters within *N. tenuis*. The final protein set from *N. tenuis* used in this analysis is available, see S1.5.

**Figure 1.**
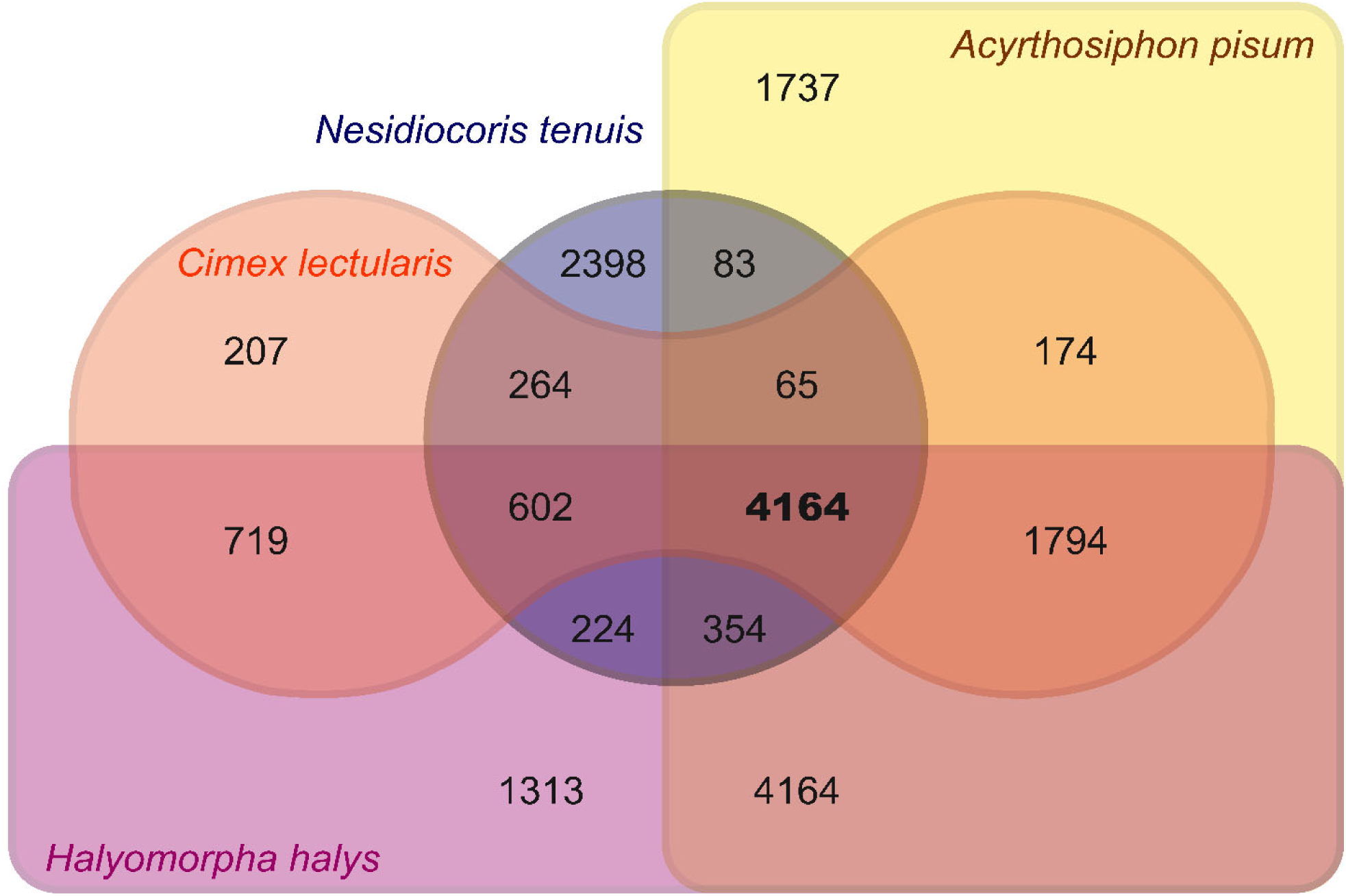
Ortholog cluster analysis of *Nesidiocoris tenuis* with three other hemipterans (*Cimex lectularis, Halyomorpha halys*, and *Acyrthosiphon pisum*). Numbers indicate the number of ortholog clusters in each grouping, with the clusters shared by all four species in bold.

**Table 3.** Output of OrthoVenn2 ortholog cluster analysis of Nesidiocoris tenuis, Cimex lectularis, Halyomorpha halys, and Acyrthosiphon pisum.

| Species | Proteins | Clusters | Singletons | Source of complete gene set |
| --- | --- | --- | --- | --- |
| *N. tenuis* | 24,668 | 8,174 | 9,136 | This work |
| *C. lectularis* | 12,699 | 7,989 | 7,989 | Poelchau *et al.*, 2015; Benoit *et al.*, 2016; Thomas *et al.*, 2020 |
| *H. halys* | 25,026 | 9,584 | 2,170 | Lee *et al.*, 2009; Poelchau *et al.*, 2015 |
| *A. pisum* | 36,195 | 8,765 | 7,298 | Legeai *et al.*, 2010; L. Xu *et al.*, 2019 |

### Karyotype analysis

Karyotype analysis revealed 2n=32 chromosomes in both females and males (Figure 2a and b). All chromosomes are relatively small with one larger pair of submetacentric chromosomes in females (Figure 2a). In males (Figure 2b), we were unable to obtain mitotic chromosomes of reasonable quality as in females and therefore we were unable to clearly identify these larger chromosomes. Screening of multiple nuclei showed sporadic deviations of the karyotype in some individuals. This was the result of supernumerary chromosomes (B chromosomes) which were clearly visible in (meiotic) pachytene stage as distinctly smaller chromosomes (Figure 2c, three B chromosomes).

**Figure 2:**
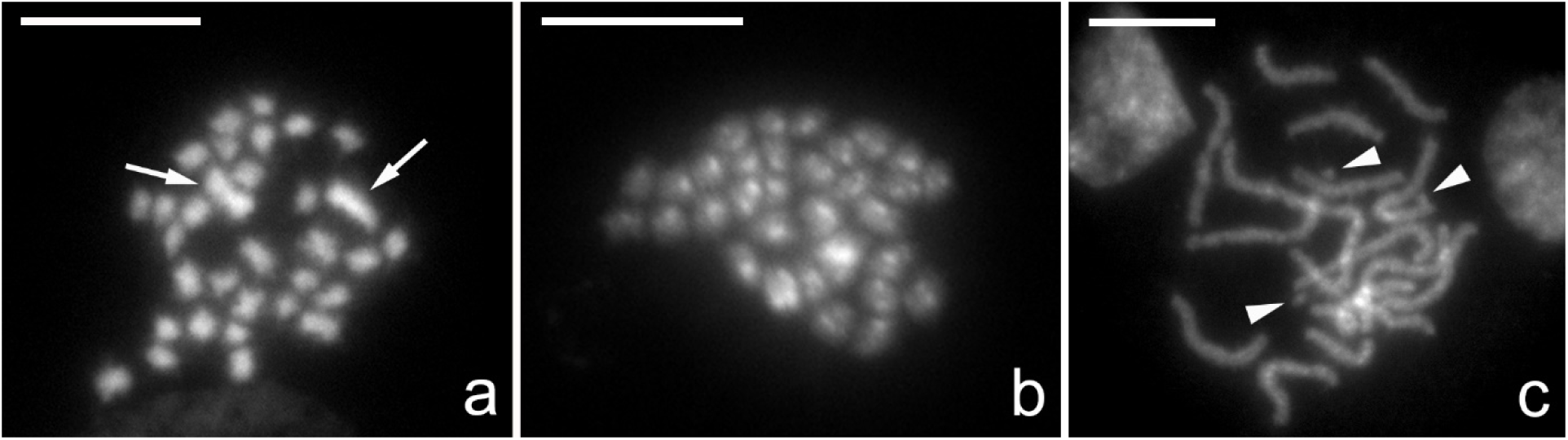
Cytogenetic analysis of *Nesidiocoris tenius* karyotype. Chromosomes were counterstained by DAPI (grey). (**a)** Female mitotic metaphase consisting of 32 chromosomes (2n=32) with two large chromosomes indicated (arrows). (**b)** Male mitotic metaphase consisting of 32 chromosomes (2n=32). (**c**) Female pachytene nucleus with B chromosomes (arrowheads). Scale bar = 10 µm.

### Analysis and localization of 18S rDNA

The 18S rDNA gene is often used as a cytogenetic marker in comparative evolutionary studies due to its ease of visualization on the chromosomes caused by high copy number and cluster organisation in animal (Sochorová et al., 2018) and plant (Gomez-Rodriguez et al., 2013) genomes. The published partial 18S sequence of *N. tenuis* (GU194646.1) and the 18S sequence identified in this study were compared to each other revealing some differences between the sequences. The published sequence consists of two fragments of 869 bp and 739 bp, which are, respectively, 99.7% and 94.2% homologous to our identified partial 18S sequence. Interestingly, the second half of our isolated 18S sequence is more homologous to a *Macrolophus sp*. partial 18S sequence (EU683153.1), i.e. 97.8%, than to the previously published *N. tenuis* sequence. A BLAST search against the *N. tenuis* genome with either of the *N. tenuis* 18S sequences resulted in four gene copies in both cases, each located on a different scaffold. However, RepeatExplorer analysis estimated 98 18S rDNA copies with the obtained genome size of 355 Mbp. Using FISH with the 18S rDNA probe we finally showed that the major rDNA forms a single cluster located terminally on a pair of homologous chromosomes (Figure 3).

**Figure 3:**
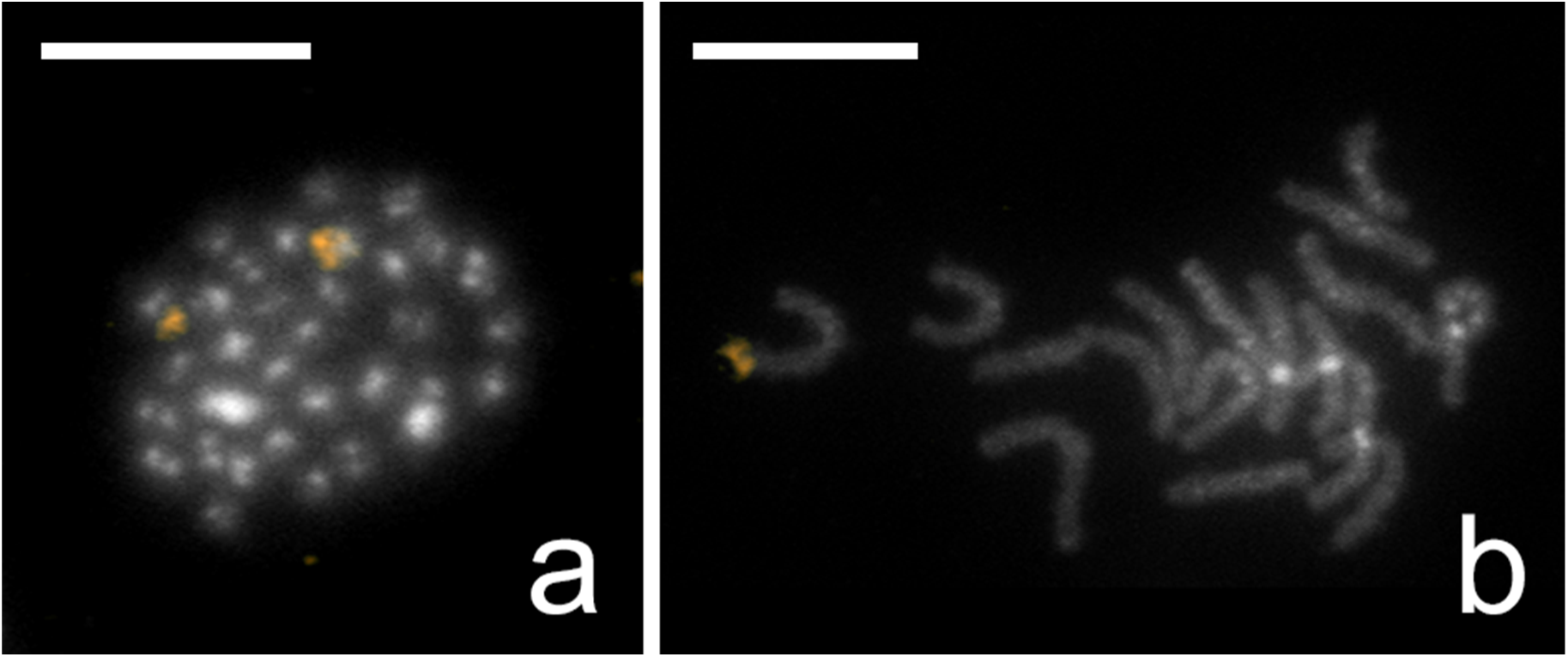
Results of *Nesidiocoris tenuis* fluorescence *in situ* hybridization with 18S rDNA probe labelled by biotin and visualised by detection with Cy3-conjugated streptavidin (gold). Chromosomes were counterstained by DAPI (grey). (**a)** Male mitotic metaphase; probe identified a cluster of 18S rDNA on two homologues chromosomes. (**b**) Female pachytene complement with one terminal cluster of 18S rDNA genes on a bivalent. Scale bar = 10 µm.

### Identification of sex chromosomes

The common sex chromosome constitution in Miridae is the male-heterogametic XX/XY system. To identify the sex chromosome constitution and estimate sex chromosome differentiation in N. *tenuis* we employed GISH and CGH experiments. The GISH results clearly revealed a single chromosome densely labelled by the male-derived probe, caused by male-enriched repetitive DNA and/or male-specific sequences which is typical for the Y chromosome (Figure 4). In addition, the Nucleolus Organizer Region (NOR; including 18S rDNA) was observed as well, as is often the case in GISH experiments due to the presence of highly repetitive sequences in the rDNA cluster. The NOR is clearly located terminally on a pair of autosomes, corroborating our 18S rDNA FISH results.

**Figure 4:**
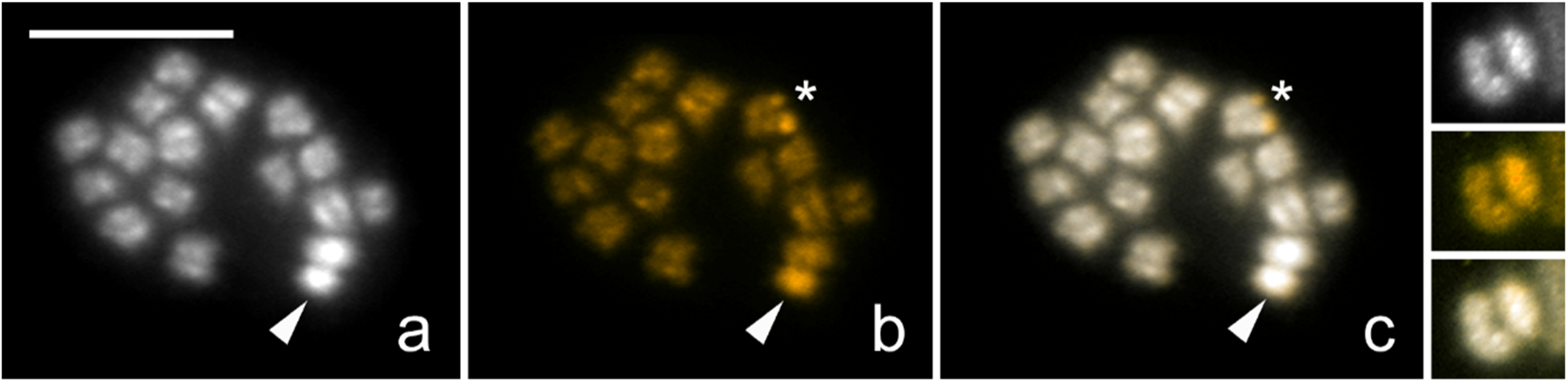
Genomic *in situ* hybridization (GISH) on male chromosomal preparation of *Nesidiocoris tenuis*. Panel (**a**) shows DAPI counterstaining (grey), panel (**b**) hybridisation signals of the male derived genomic probe labelled by Cy3 (gold) together with competitor generated from unlabelled female genomic DNA, and panel (**c**) a merged image. (**a**, **b**, **c**, **detail**) Meiotic metaphase I, male derived probe highlighted the Y chromosome (arrowhead) more (**b**, **c**) compared to autosomes and the X chromosome. Note highlighted terminal regions of one of the bivalents caused by presence of major rDNA genes (asterisk). (**detail**) Detail picture of XY bivalent; Y chromosome labelled by male derived probe. Note that the Y chromosome is smaller in size and showing more heterochromatin compared to the X chromosome (and autosomes). Scale bar = 10 µm.

To further study the differentiation of the sex chromosomes we carried out CGH experiments on chromosome preparations of both sexes (Figure 5). All chromosomes were labeled evenly by the female and male probes with the exception of the largest chromosome pair. Both sex chromosomes were highlighted with DAPI (Figure 5a, e), indicating that they are both A-T rich and largely composed of heterochromatin. In females, the largest chromosome pair was labelled more by the female probe than the male probe indicating that these chromosomes contain sequences with higher copy numbers in females, and are thus the X chromosomes, as seen in Figure 5a-d. In male meiotic nuclei (Figure 5e-h), two types of nuclei can be discerned, where the largest chromosome was labelled more by either the female probe or the male probe corresponding to the X, and Y chromosome, respectively, whereas the autosomes were labelled equally by both probes.

**Figure 5:**
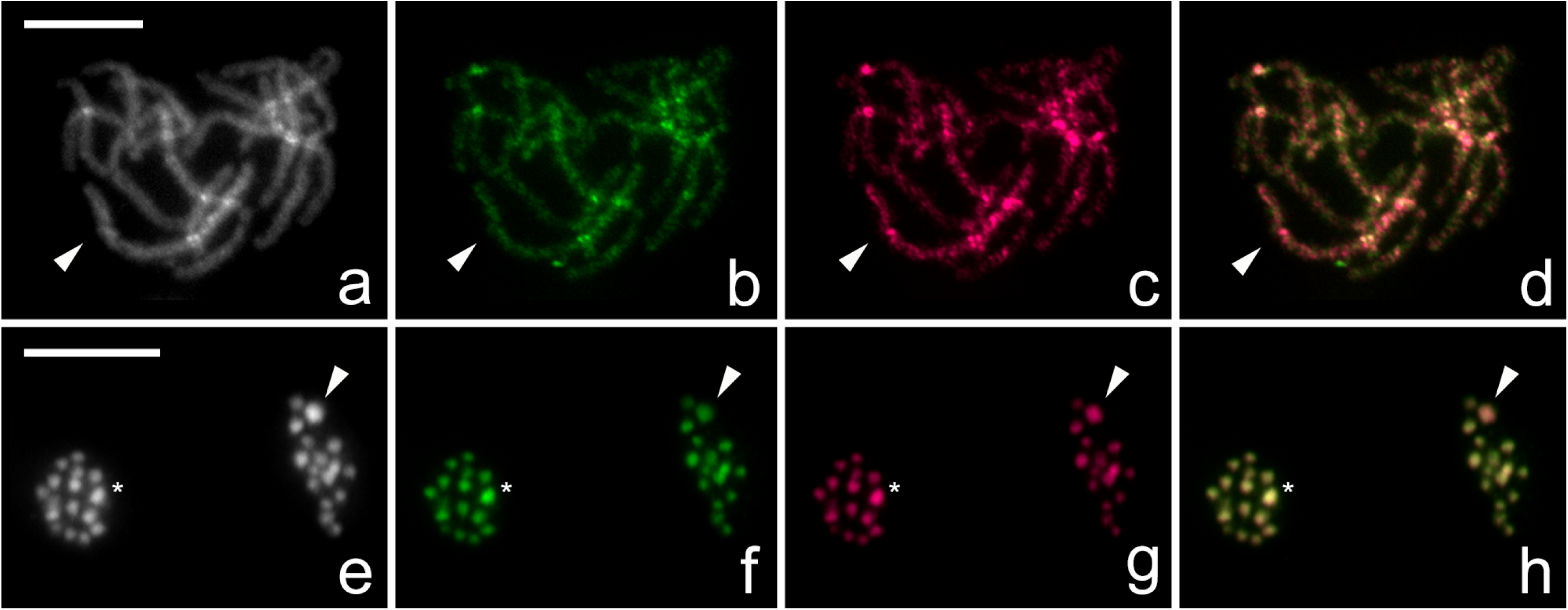
Comparative genomic hybridization (CGH) on female (**a**, **b**, **c**, **d**) and male meiotic metaphase II (**e**, **f**, **g**, **h**) chromosomes of *Nesidiocoris tenuis*. Panels (**a**, **e**) show chromosomes counterstained by DAPI (grey), panels (**b**, **f**) hybridization signals of the male derived genomic probe labelled by fluorescein (blue), panels (**c**, **g**) hybridization signals of the female derived genomic probe labelled by Cy3 (gold), and panels (**d**, **h**) merged images. (**c**, **d**) Note that the X chromosome bivalent (arrowhead) in female pachytene complement was highlighted more by female probe compared to the autosomal bivalents; (**b**, **d**) male probe labelled all chromosomes equally. (**h**) Two sister nuclei in meiotic metaphase II showed equal hybridization patterns of both probes on autosomes; in one of the forming nuclei, the X chromosome (arrowhead) was highlighted by female derived genomic probe (**g**, **h**) and in the second nucleus the Y chromosome (asterisk) was strongly highlighted by male derived genomic probe compared to autosomes (**f**, **h**) and less highlighted by female derived probe (**g**, **h**). (**e**) Note that the sex chromosomes are the biggest and most heterochromatic elements in the nucleus. Scale bar = 10 µm. **Note: Alternate colouration available in supplementary material (S1.6.3).**

### Identification and mapping of abundant repeats

RepeatExplorer software was used on reads with GEM barcodes removed to identify the most abundant repeats in the genome of *N. tenuis* (results available in supplementary materials, see Table S1.6.1). The most abundant repeat, Nt_rep1, makes up approximately 3% of the genome estimated by RepeatExplorer. Analysis on the assembled genome, using a coverage cut-off value of 70%, reveals that Nt_rep1 is present on 3190 scaffolds (8.737% of the assembled scaffolds), with a maximum of 17 copies on a single scaffold. According to the assembled genome, Nt_rep1 makes up approximately 0.8% of the entire genome (Figure S1.6.2). We subsequently mapped Nt_rep1 to the chromosomes of *N. tenuis* using FISH. The repeat is located on most chromosomes and is accumulated in sub-telomeric regions (Figure 6). Additional signals were identified on the X chromosome indicating a higher number of this repeat (Figure 6a-c). This increase in frequency is specific to the X chromosome and is not found on the Y chromosome of *N. tenuis* (Figure 6d-f).

**Figure 6:**
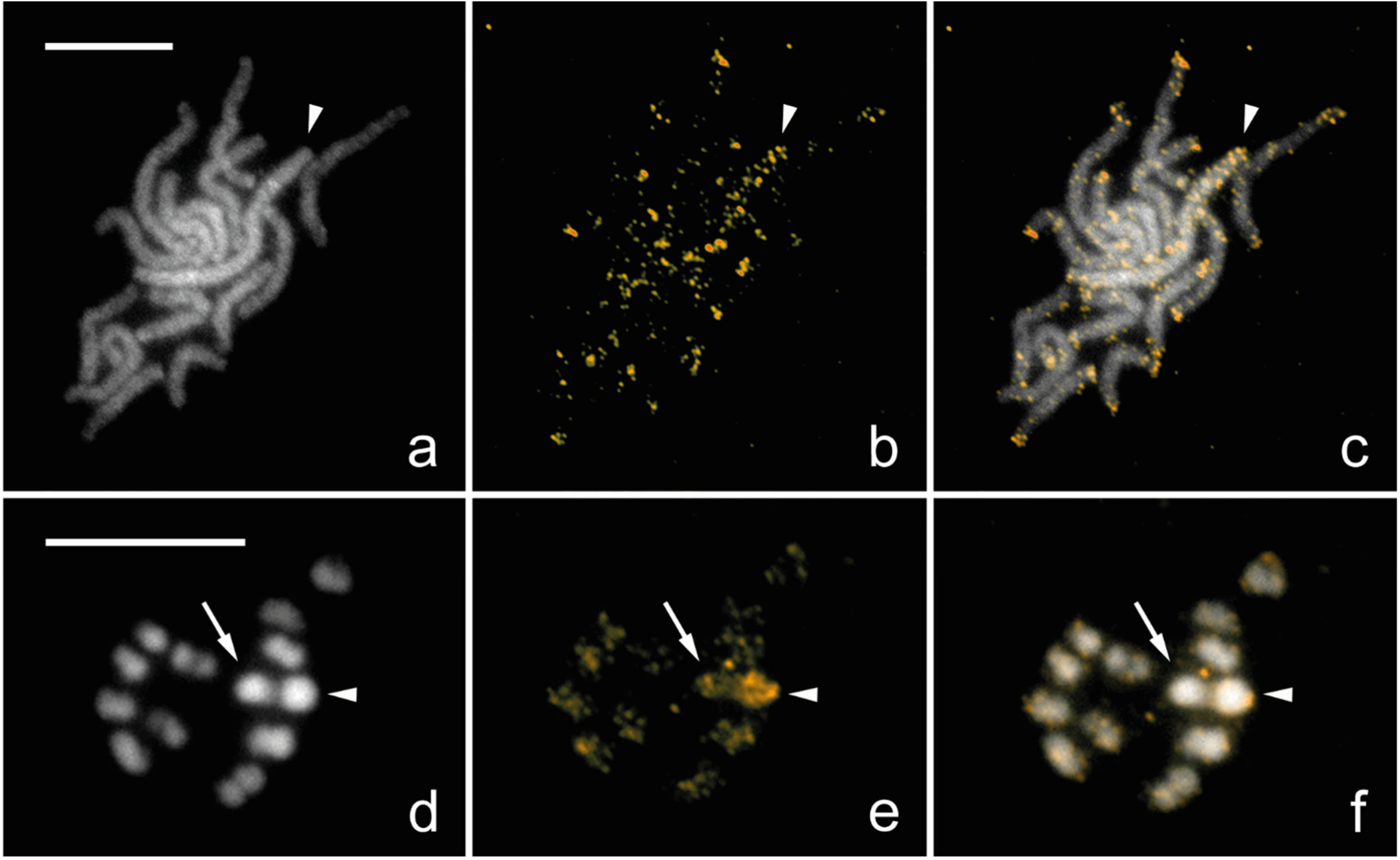
Fluorescence *in situ* hybridization with Nt_rep1 probe labelled by biotin (gold) on female (**a**, **b**, **c**) and male (**d**, **e**, **f**) chromosomes of *Nesidiocoris tenuis* counterstained with DAPI (grey). (**a**, **b**, **c**) Female pachytene chromosomes; Nt_rep1 probe highlighted the pair of X chromosomes (arrowhead). (**b**, **c**) Note that terminal regions of all bivalents were also labelled by probe, probably due to presence of this sequence in sub-telomeric regions. (**d**, **e**, **f**) Incomplete male nucleus in meiotic metaphase I; probe highlighted the X chromosome (arrowhead) more compared to autosomes and Y chromosome (arrow). (**b**, **c**, **e**, **f**) Strong hybridization signals on X chromosomes in both sexes were caused by enrichment of Nt_rep1 sequence on the X chromosomes. Scale bar = 10

### Testing of candidate telomere motifs

Analysis of the raw sequencing data and the assembled genome both revealed low numbers of the insect telomere motif (TTAGG)*_n_* (Frydrychová et al., 2004) in *N. tenuis*, i.e. approximately 98 repeats per haploid genome. This translates into approximately three copies of the repeat per chromosome end, much lower than expected for a telomeric motif. These low copy numbers were additionally confirmed using Southern dot blot (Figure S1.6.1). Other candidate telomere motifs previously identified by TRF analysis, (TATGG)*_n_*, (TTGGG)*_n_*, and (TCAGG)*_n_*, were examined by FISH for their distribution in the genome. They were found scattered throughout the genome but lacked a clear accumulation at the terminal regions of the chromosomes (not shown). Therefore, these sequences can also be excluded as telomeric motifs in *N. tenuis*.

### Pooled sequencing analysis

Using previously generated whole genome sequencing of ten females from the KBS population, we were able to estimate genetic diversity of the commercial population via a pool-seq population analysis. Read coverage was randomly subsampled to 10X coverage (18,000,000 reads). Additionally, we used a modified v1.5 *N. tenuis* genome with scaffolds of less than 10,000 bp removed. This was to ensure that window sliding was not being inflated on scaffolds smaller than the window size. This reduced the genome from 36,513 scaffolds to 7,076, however, the reduced genome still contained 72.23% of the genome in terms of size (256,487,768 bp).

Three runs of PoPoolation were performed with varied window size, step size, and the masking of indel regions. The default setting, window size and step size of 10,000 bp, yielded similar results as the adjusted window size and step size of 5,000 bp, while differences were apparent when indel regions were masked. As such, results of window size and step size 10,000 bp with indel regions mapped are reported here (other results available in S1.7). The variance sliding program created 28,833 windows of 10,000 bp with mapped reads, of which 5,913 were sufficiently covered with reads to calculate values per window (coverage ≥ 0.60). Genome-wide, the nucleotide diversity (*π*) is 0.0080 and *Tajima’s D* is −0.0355. Figure 7 shows the *Tajima’s D* (7a) and *π* (7b) for the ten largest scaffolds, all containing gene annotations, arranged in order of size. These ten scaffolds represent approximately 1.7% of the genome (6,135,756 bp), and varied in terms of window coverage (from no coverage to full coverage) as well as both *Tajima’s D* and *π*. These ten scaffolds are a snapshot of the whole genome, summarised in Table 4, whereas genome-wide results can be found in S1.7.

**Figure 7.**
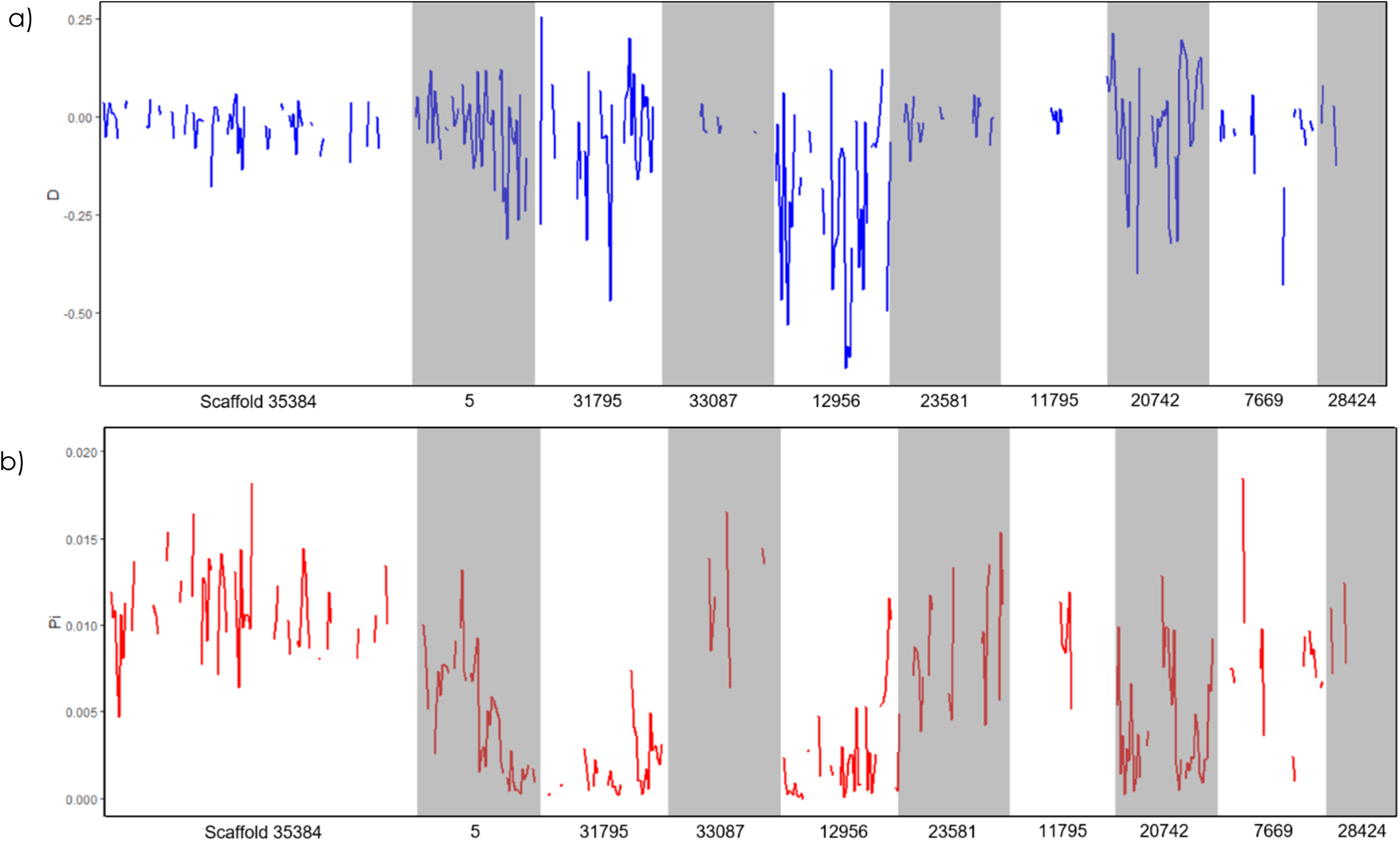
Genetic diversity of Koppert Biological Systems commercial *Nesidiocoris tenuis* population according to *Tajima’s D* (**a**), and nucleotide diversity, *π* (**b**). Scaffolds are ordered according to size, which each name beneath on the x-axis.

**Table 4.**
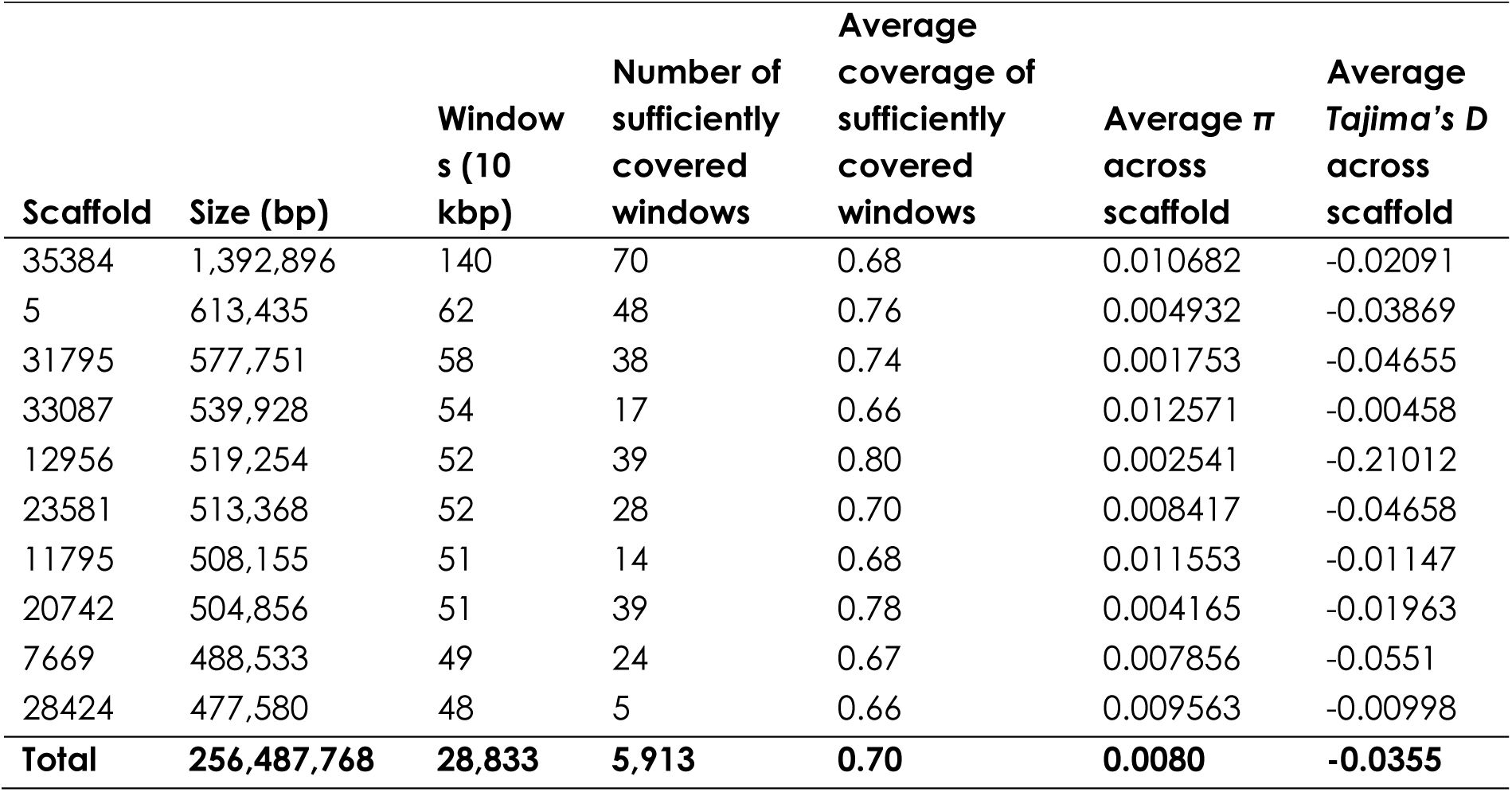
PoPoolation analysis on commercial Koppert Biological Systems Nesidiocoris tenuis population (n=10 females), with 10 largest scaffolds according to size. Coverage is ≥ 0.60 and indel regions are masked.

| **Scaffold** | **Size (bp)** | **Windows (10 kbp)** | **Number of sufficiently covered windows** | **Average coverage of sufficiently covered windows** | **Average $\pi$ across scaffold** | **Average Tajima's D across scaffold** |
| --- | --- | --- | --- | --- | --- | --- |
| 35384 | 1,392,896 | 140 | 70 | 0.68 | 0.010682 | -0.02091 |
| 5 | 613,435 | 62 | 48 | 0.76 | 0.004932 | -0.03869 |
| 31795 | 577,751 | 58 | 38 | 0.74 | 0.001753 | -0.04655 |
| 33087 | 539,928 | 54 | 17 | 0.66 | 0.012571 | -0.00458 |
| 12956 | 519,254 | 52 | 39 | 0.80 | 0.002541 | -0.21012 |
| 23581 | 513,368 | 52 | 28 | 0.70 | 0.008417 | -0.04658 |
| 11795 | 508,155 | 51 | 14 | 0.68 | 0.011553 | -0.01147 |
| 20742 | 504,856 | 51 | 39 | 0.78 | 0.004165 | -0.01963 |
| 7669 | 488,533 | 49 | 24 | 0.67 | 0.007856 | -0.0551 |
| 28424 | 477,580 | 48 | 5 | 0.66 | 0.009563 | -0.00998 |
| **Total** | **256,487,768** | **28,833** | **5,913** | **0.70** | **0.0080** | **-0.0355** |

## Discussion

### Assembly and Annotation

Presented here is the genome of *N. tenuis,* a biological control agent used throughout the Mediterranean in tomato crops. We chose to use 10X Genomics linked-read sequencing strategy as it best suited the challenges that come with working with a relatively small and long-lived mirid such as *N. tenuis.* Assembling a genome is easiest with reduced heterozygosity in the input sample, often through single individual sampling or inbreeding (Ekblom and Wolf, 2014; Richards and Murali, 2015). This proved an initial challenge for the sequencing strategy of *N. tenuis*, as they are too small for a single individual to yield the minimum amount of DNA required for a traditional NGS library, and an inbred population was not readily available for sequencing. Therefore, 10X Genomics linked-read sequencing was the immediate solution for which a small amount of input DNA from a single individual would yield a highly contiguous genome.

Assemblies v1.0 and v1.5 contain 5.91% and 6.29% ambiguous nucleotides, while still offering a relatively high BUSCO score, with the final decontaminated assembly (v1.5) having a completeness of 87.5% of the insect_0db9 ortholog dataset. However, the final assembled genome size is approximately 150 Mbp larger than was expected based on flow cytometry data, and we suggest the assembly presented here can best be improved in terms of accuracy and contiguity with long reads from an inbred sample. This discrepancy between estimated genome size and assembled genome size may also be due to the ambiguous nucleotides inserted into the genome during the assembly process. Making up just over 6% of the final assembled genome, that is approximately 2.2 Mbp of ambiguous nucleotides. However, most of the genome inflation is likely due to residual contamination along with duplicate scaffolds that remain after removing 100% identical ones.

Annotation via evidence-based, homology-based, and *ab initio* models resulted in 24,668 genes. Compared to other assemblies within the hemipteran order, such as *C. lectularis*, with a genome size of 650 Mbp and 12,699 genes (Thomas et al., 2020) or *A. pisum*, with a draft genome size of 464 Mbp and 36,195 genes (Richards et al., 2010), *N. tenuis* sits, in the middle in terms of genome size and number of genes. It is worth noting that of the 24,668 genes within the complete gene set, only 11,261 (45.7%) remained after UniProtKB mapping, of which 8,920 (36.2% of total) were used by DAVID for functional analysis. This is relatively low compared to similar genome projects, such as *Aphys gossypii* (Glover) (Hemiptera: Aphididae), where 49.2% of the gene set could be used for GO term analysis (Quan et al., 2019). However, we used different methods of gene prediction and annotation which may explain the difference. The next step for the *N. tenuis* genome is manual annotation and curation, which would likely improve the GO term analysis, but this requires time and expertise. Still, we hope that other researchers will use and add to the annotation.

Comparing the current gene set of *N. tenuis* to other Hemipterans, the clustering identified considerable overlap, as 71% of the clusters that are found in *N. tenuis* were shared between the other species in the comparative analysis. Despite being more closely related to *C. lectularis* in terms of phylogeny, in terms of lifestyle, *N. tenuis* is far more similar to *A. pisum* and *H. halys*, and this is likely reflected in absolute number of proteins and clusters shared between the four species. The remaining 29% of clusters, as well as the singleton proteins, are indications for proteins unique to either *N. tenuis* or Miridae in general. Through the OrthoVenn2 website, the analysis performed here can be easily replicated, altered with other species of interest, and even improved upon if the complete gene sets are updated or with a newer software version. In our iteration, the 24,668 proteins of *N. tenuis* group into 8,174 clusters. 2,398 clusters are unique to *N. tenuis*, however, some of these genes have a relatively strong homology to genes of one of the other species used in the analysis and could be incorrectly flagged as being unique. Reasons could be poor gene annotation quality resulting in a poor *in silico* protein translation, or too stringent clustering settings. Regardless, these 2,398 clusters may be of interest to researchers working on zoophytophagy, the negative effects of *N. tenuis* on tomato as compared to other mirids, or broader questions such as phylogeny of the Hemiptera.

### Characterizing the Genome

Every sequence and assembly strategy has benefits and drawbacks, and the 10X Genomics linked-read strategy is no exception. The technique requires only few nanograms of DNA for library preparation which allowed us to use a single individual and removed the need for inbreeding to reduce variation in the sequencing population. However, using a single individual from a closed and proprietary rearing process presented other challenges. These challenges were threefold: we had to deal with bacterial contamination as antibiotic treatment is not possible, we had to ensure that the single individual-derived assembly reflects reality in terms of genes present and structure, and we had to ensure that a single female-derived assembly is applicable for population-level analyses.

Contamination of genomes is a constant concern, and sequencing strategies should attempt to address the risks in the best way possible to deliver reliable genomes (Ekblom and Wolf, 2014). Equally so is the desire for inbred strains if multiple individuals are required to reach the micrograms of DNA necessary for NGS platforms. The inability to remove symbionts and microbiota using antibiotics administered to a few successive generations as well as the difficulty or inability to inbreed a strain is not restricted to *N. tenuis*. The sequencing strategy chosen for the mountain pine beetle, *Dendroctonus ponderosae* (Hopkins) (Coleoptera: Curculionidae), relied on assuming the relatedness of several individuals as well as isolating the gut during the extraction process, and still additional post-assembly decontamination was required (Keeling et al., 2013). A linked-read strategy with low input requirements, such as the 10X Genomics library preparation that was chosen here, negates the need for a pool of inbred samples or controlling for relatedness. However, another potential benefit of controlled rearing such as those used in inbreeding (as opposed to be limited to wild-caught specimens, for example) is the ability to treat with antibiotics for multiple generations. Without the ability to do so, sequencing and assembly strategies rely heavily on post-sequencing decontamination strategies (both pre- and post-assembly are possible). That such post-assembly filtering strategies as used here for *N. tenuis* can be successful was shown by a less contaminated assembly and by the identification of potential LGTs.

### Beyond the Genome: Potential symbionts and LGT events

The list of potential symbionts or pathogens generated in Table 2 represent both insect and plant pathogens, as well as potential environmental contaminants. In addition to the positive test for *Wolbachia* in the KBS population used here, *N. tenuis* is known to potentially harbour *Rickettsia* as an endosymbiont in addition to *Wolbachia* (Caspi-Fluger et al., 2014). *Rickettsia* genome sizes can range from 0.8 to 2.3 Mbp, also reflecting variation in levels of reductive evolution (Sachman-Ruiz and Quiroz-Castañeda, 2018). However, the total scaffold length identified here as from a Rickettsia falls below this range, likely indicating incomplete recovery of the genome from the insect sequencing. The potential symbionts revealed included not only *Wolbachia* and *Rickettsia*, but also other known insect symbionts, in addition to the usual lab contamination suspects.

*Sodalis* is a genus of bacterium symbiotic with various insects, including the tsetse fly and louse fly, louse and hemipteran species (Boyd et al., 2016). Genome sizes of *Sodalis* and close relatives range from 0.35 to 4.57 Mbp (Santos-Garcia et al., 2017). The relatively small total scaffold size found in our results (0.36 Mbp) likely reflects incomplete genome recovery in the assembly, but could also be due to genome size reduction, and is worthy of further investigation. *Erwinia* and *Pantoea* are closely related bacteria that are associated with plant pathology (Kamber et al., 2012; Zhang and Qiu, 2015) and both have been found in the midgut of stink bugs as vertically transferred plant-associated bacteria that become temporary endosymbionts of stink bugs until later replacement with another endosymbiont (Prado and Almeida, 2009). The genome sizes of *Erwinia* and *Pantoea* species typically range from 3.8-5.1 Mb. Our total scaffold size for the *Erwinia* and *Pantoea* are substantially smaller (2.078 Mbp and 2.22 Mbp), but it is possible that these scaffolds belong to the same bacterium. In any case, the association of a zoophytophagous mirid bug with potential plant pathogens is noteworthy, especially in a biological control context.

As for *Serratia*, *Serratia marcescens* is both a common Gram-negative human-borne pathogen and a causal agent of cucurbit yellow vine disease (CYVD) (Abreo and Altier, 2019; Bruton et al., 2007). It is worth noting that in cases of CYVD, the transmission of *Serratia marcescens* from its vector the squash bug, *Anasa tristis* (De Geer) (Hemiptera: Coreidae), to host crops is via the phloem. Other *Serratia* spp. have been identified as insect symbionts previously, as have other potential symbionts found in the contaminated scaffolds, such as *Cedecea* spp. (Jang and Nishijima, 1990). The presence of *Dickeya* is an interesting find, as *Dickeya dadantii* has been established as a pathogen of *A. pisum*, while the pea aphid itself is a potential vector for the bacterium with regards to plants (Costechareyre et al., 2012). *Dickeya* spp. cause soft rot in various crops, including tomato. In a similar vein, the identification of *Ralstonia,* as some members of this order, such as *Ralstonia solanacearum,* are soil-borne pathogens that causes wilt in several crop plants, including tomato (Lowe-Power et al., 2018). However, both *Dickeya* spp. and *R. solanacearum* infect the xylem while *N. tenuis* is a phloem-feeder. The scaffold lengths for *Citrobacter* and *Cronobacter* are also considerably below their typical genome sizes, and likely represent incomplete sequence recovery in the metagenomic sample.

All of these described associations are preliminary, as follow-up analyses against the entire NCBI database and proper bacterial gene annotation are lacking. Nevertheless, these putative bacterial associations of *N. tenuis*, their distribution within the insect, and their possible biological significance, warrants further investigation. It is important to note that some of these bacterial “contaminants” may actually represent large LGTs, which can be confirmed by identifying flanking sequences (e.g. using long-read technologies) and/or *in situ* chromosome hybridization analyses, such as done for the large LGT in *Drosophila ananassae* (Doleschall) (Diptera: Drosophilidae) (Dunning Hotopp et al., 2007). More research into the symbionts of *N. tenuis* via metagenomics would certainly shed some light on true symbionts (or pathogens) versus true contaminants, with potential implications for biological control and related research.

One of the LGT candidate genes that were detected following manual curation corresponds to phenazine biosynthesis protein *PhzF* (OIV46256.1), with the likely microbial source being a *Sodalis* species. Phenazines are heterocyclic metabolites with “antibiotic, antitumor, and antiparasitic activity,” but are also toxic when excreted by bacteria (Blankenfeldt et al., 2004). This LGT region exhibits gene expression and is flanked by conserved insect genes, providing further support for it being a legitimate LGT, though further research into this region will be necessary to confirm this. The gene occurs on two different scaffolds, 22012 and 22013, which are highly similar to each other in some regions at the nucleotide level. These could represent homologous regions that differ sufficiently to assemble as different scaffolds, or alternatively a duplication in two different regions of the genome. Future work should focus on its expression patterns in different tissues (e.g. salivary glands, in interest of *PhzF*) and potential functional role in *N. tenuis*.

### Beyond the Genome: Cytogenetics

We determined the karyotype of *N. tenuis* to be 2n=32 (30+XY in males) chromosomes which is the second most common chromosome number in the family Miridae (Kuznetsova et al., 2011). In addition, we have shown that *N. tenuis* has an XX/XY sex chromosome constitution, with the sex chromosomes being the largest elements in the karyotype. This is different from the closely related *Macrolophus costalis* (Fieber) (Hemiptera: Miridae) (2n=24+X1X2Y), and *M. pygmaeus* (Rambur) (2n=26+XY) where two pairs of autosomes are larger than the sex chromosomes, yet similar to *M. melanotoma* (Costa) which only differs from *N. tenuis* in the number of autosomes, 2n=32+XY (Jauset et al., 2015). As we sequenced a single female, sequence information of the Y chromosome is missing from our genome assembly. While analyzing the *N. tenuis* karyotype we discovered the sporadic presence of B chromosomes in the KBS population. B chromosomes are supernumerary chromosomes that are dispensable to the organism, and are often present in only a subset of individuals from a population (Banaei-Moghaddam et al., 2013). Supernumerary chromosomes are common in Heteroptera, yet only a few species of Miridae have been identified to carry supernumerary chromosomes (Grozeva et al., 2011). Presence of B chromosomes in high numbers within an individual is often found to be detrimental, though in lower numbers they are often considered neutral or, in some cases, beneficial (Camacho et al., 2000; Jones and Rees, 1982). The abundance of B chromosomes in *N. tenuis* biological control populations is currently unknown but determining their potential effects on fitness-relevant traits might reveal beneficial information for the optimization of mass-reared populations.

The hemizygous sex chromosomes of most organisms have a high content of repetitive DNA, consisting of multiple different repetitive sequences that are less frequent found on autosomes (Charlesworth and Charlesworth, 2000; Traut et al., 1999). Therefore, the use of cytogenetic techniques, such as CGH and GISH, in the identification of hemizygous sex chromosomes is a powerful tool and is well established in different groups of organisms, e.g. Lepidoptera (Carabajal Paladino et al., 2019; Dalíková et al., 2017a; Zrzavá et al., 2018), Orthoptera (Jetybayev et al., 2017), fish (Sember et al., 2018), and frogs (Gatto et al., 2018). However, to our knowledge, this is the first time these techniques have been used in the family Miridae. The X and Y chromosome of *N. tenuis* are similar in size, with the X chromosome being slightly bigger, and are difficult to distinguish from each other based solely on their appearance without special probing. Our CGH and GISH results showed relatively weak hybridization signals of genomic probes on the sex chromosomes compared to other species indicating little differentiation of sequence content between the X and Y chromosomes, and/or between the sex chromosomes and the autosomes. Though the hybridization signals are relatively weak, not only the Y chromosome but also the homogametic sex chromosome, the X chromosome, is distinguishable in the CGH results, which shows X-enriched or X-specific repetitive DNA, similar to what was found on the Z chromosome in *Abraxas* spp. (Zrzavá et al., 2018). Mapping of the most abundant repeat in the genome revealed that one such X-enriched repeats is Nt_rep1, confirming the outcomes of our CGH results.

The low copy numbers of 18S rDNA identified in the assembled genome were surprising. The NOR is usually composed of tens to hundreds of copies, and is therefore used in heteropteran cytogenetic studies due to its easy visualization (Kuznetsova et al., 2011). Analysis of the raw data estimates 98 copies of 18S rDNA are present in the genome, yet the majority of these copies are missing from the final assembly. The FISH results show that 18S rDNA is present as a single cluster in the genome, indicating that there is a limit to the genome assembler Supernova, and 10X Genomics by extension, and its ability to assemble highly repetitive regions of the genome.

Similarly, the FISH results of Nt_rep1 and the analysis of the copy numbers and distribution of the repeat in the genome assembly do not corroborate. Though many copies of the repeat are present in the assembled genome, most scaffolds contain one or few copies of the repeat. The FISH results, however, show multiple clusters scattered across most chromosomes each containing high copy numbers, revealing a lack of scaffolds containing high copy numbers of Nt_rep1 in the assembled genome. Therefore, analyses on repetitive DNA content are currently more reliable using the short sequence reads rather than the assembled genome as it underestimates repeat content. Long read sequencing methods would be able to overcome such problems with repetitive DNA, not only in *N. tenuis* but in any species, and would be better suited to analyse repetitive regions of genomes. As mentioned before, a hybrid assembly strategy combining our 10X sequencing data with long reads, obtained by e.g. Oxford Nanopore or PacBio sequencing, would presumably improve the assembly, though in this aspect for particular segments of the genome that are high in repetitive DNA. This should be kept in mind for other 10X Genomics/Supernova-derived genomes: the true number of repeats may be underestimated.

Screening of the genome and Southern blot assay suggests the absence of the ancestral insect telomere motif, (TTAGG)_n_, *in N. tenuis*, as the case in other species from the family Miridae (Grozeva et al., 2019; Kuznetsova et al., 2011). The telomeric motif was present in our Tandem Repeat Finder results, but in much lower numbers than expected for telomeric sequences. Additional attempts of identifying the telomeric repeat motif did not resolve this question. Three additional repeats we identified in the *N. tenuis* genome were tested via FISH, i.e. (TATGG)_n_, (TTGGG)_n_, and (TCAGG)_n_, but did not localise near the ends of the chromosomes. Notably though, mapping the most abundant repeat in the genome, Nt_rep1, did reveal accumulation in the sub-telomeric regions of chromosomes (Figure 6). Therefore, our approach to identify potential telomere motifs, though presently unsuccessful, would presumably be effective if more repeats would be screened. In addition, a similar approach was used by Pita et al. (2016) in *T. infestans*, where the insect telomere motif, (TTAGG)_n_, was successfully identified from the raw sequencing data (Pita et al., 2016). It must be noted, however, that the telomeres of *N. tenuis* might consist of different types of repeats other than short tandem repeats (as found in, for example, *Drosophila*; Traverse & Pardue, 1988) which would not be identified using Tandem Repeat Finder (Traverse and Pardue, 1988). Therefore, the identity (or even presence) of the telomeric repeat in *N. tenuis,* and by extension Miridae, remains unknown.

### Beyond the Genome: Population Genomics

Pooled sequence data of ten females from the KBS population were compared against the genome and provide interesting population-level effects. The overall negative *Tajima’s D* would seem to indicate an abundance of rare alleles and is possible evidence of selective sweeps or population expansion, as seen in some populations *of Drosophila serrata* (Malloch) (Reddiex et al., 2018), however, this generally results in more negative values (near −1 or −2). While overall negative, the absolute value of *D* in our results is small in comparison (total range from −0.89 to 0.56). To best assess the state of the commercial population, monitoring the genetic variation over time would indicate if the population is undergoing an expansion after a bottleneck (*D* < 0) or contracting (*D* > 0), whereas when *D* = 0, we assume no selection. We can then assume that there is no selection currently at play in the commercial population. The few studies that have looked at genetic diversity within biological control populations have primarily been reduced representation analyses, such by genotyping with microsatellites (Paspati et al., 2019). Here, a pool-seq approach offers a genome-wide look at the population and can give indications of the genetic diversity of the population; this could be a useful tool for monitoring population levels efficiently and determining which regions of the genome are under selection in a biological control context.

Both genetic diversity values calculated here can also be used in population comparisons between the biological control population and wild populations. For instance, Xun *et al*. used mitochondrial and nuclear barcoding regions to haplotype 516 individuals across 37 populations into two regional groups, southwest China (SWC) and other regions in China (OC) (Xun et al., 2016). *π* was 0.0048 (SWC) and 0.904 (OC), while *D* was −0.112 (SWC) and −1.998 (OC). It was concluded that the SWC population was stable while, similar to the KBS population here, the OC population was undergoing sudden population explosion. Pooled sequencing could be a useful tool for comparing wild Mediterranean populations to the commercial population to determine disparities in genetic variation as well as to understand the dynamics of the wild populations.

There is a concern in using PoPoolation in this context: are ten individual females sufficient for determining population variation? Here we used existing population sequence data to better utilize resources, reduced to an appropriate coverage with masked indel regions. This enabled us to show population-level impacts at the very least, which can then pave the way for further studies, with better constructed sampling methods and sample sizes; the lack of perfect data should not preclude preliminary studies from being pursued.

## Conclusion

Reported here is the genome for *N. tenuis*, a mirid that is both used throughout the Mediterranean Basin as a biological control agent and reported as a greenhouse pest in other European countries. The assembled genome is 355 Mbp in length, composed of 42,830 scaffolds with an N50 of 27,074 bp. The goal of this project was to not only provide a genome, but also to highlight possible avenues of research now available with *N. tenuis*. A protein analysis has provided interesting prospects for mirid-specific proteins, while examples of potential LGT call for further inquiry. Putative symbionts were identified while filtering out contamination, creating a precursor for future metagenomic analysis. The cytogenetic analyses of *N. tenuis* here shed some light on Mirid cytogenetics, such as the karyotype and sex determination system, but also solicits more questions. As for the commercial population, now that there is a baseline level of genetic variation documented through our pooled sequencing, what remains to be seen is how it compares to other populations, such as other commercial populations, wild, or invasive populations. To this end, future exploration on these themes, among others, are now greatly facilitated with our release of this genome.

## Acknowledgements

We would like to express special thanks to Milena Chinchilla Ramírez for her advice and *N. tenuis* expertise. Thanks to Markus Knapp (Koppert BV), Javier Calvo (Koppert BS) and Gerben Messelink (WUR) for providing *N. tenuis* specimens. Thanks to both José van de Belt, Frank Becker (WUR), and Carolina Gallego (IVIA) for their technical assistance in DNA and RNA extraction. We acknowledge the assistance of Magda Zrzavá, Anna Voleníková, Martina Flegrová, and Diogo Cabral-de-Mello (Institute of Entomology BC CAS) for their cytogenetic expertise and assistance. JHW acknowledges Sammy Cheng for assistance with the LGT pipeline. KBF acknowledges Jetske de Boer for her assistance with the flow cytometry and Joost van den Heuvel for his assistance with PoPoolation. Annotation was performed by GenomeScan B. V (NL). Access to computing and storage facilities owned by parties and projects contributing to the National Grid Infrastructure MetaCentrum (CZ), provided under the programme “Projects of Large Research, Development, and Innovations Infrastructures” (CESNET LM2015042), is greatly appreciated. This project was funded by the European Union’s Horizon 2020 research and innovation program under the Marie Skłodowska-Curie grant agreement no. 641456. JHW acknowledges the US-NSF-IOS-1456233 and Nathaniel & Helen Wisch Chair for funding support. The research leading to these results was partially funded by the Spanish Ministry of Economy and Competitiveness MINECO (RTA2017-00073-00-00). Cytogenetic experiments were financed by grants 17-13713S (SV and FM) and 17-17211S (MD and IP) of the Czech Science Foundation.

## References

Abreo, E., Altier, N., 2019. Pangenome of Serratia marcescens strains from nosocomial and environmental origins reveals different populations and the links between them. Sci. Rep. 9, 1–8. https://doi.org/10.1038/s41598-018-37118-0

Acland, A., Agarwala, R., Barrett, T., Beck, J., Benson, D.A., Bollin, C., Bolton, E., Bryant, S.H., Canese, K., Church, D.M., Clark, K., Dicuccio, M., Dondoshansky, I., Federhen, S., Feolo, M., Geer, L.Y., Gorelenkov, V., Hoeppner, M., Johnson, M., Kelly, C., Khotomlianski, V., Kimchi, A., Kimelman, M., Kitts, P., Krasnov, S., Kuznetsov, A., Landsman, D., Lipman, D.J., Lu, Z., Madden, T.L., Madej, T., Maglott, D.R., Marchler-Bauer, A., Karsch-Mizrachi, I., Murphy, T., Ostell, J., O’Sullivan, C., Panchenko, A., Phan, L., Pruitt, D.P.K.D., Rubinstein, W., Sayers, E.W., Schneider, V., Schuler, G.D., Sequeira, E., Sherry, S.T., Shumway, M., Sirotkin, K., Siyan, K., Slotta, D., Soboleva, A., Soussov, V., Starchenko, G., Tatusova, T.A., Trawick, B.W., Vakatov, D., Wang, Y., Ward, M., John Wilbur, W., Yaschenko, E., Zbicz, K., 2014. Database resources of the National Center for Biotechnology Information. Nucleic Acids Res. 42, 8–13. https://doi.org/10.1093/nar/gkt1146

Andrews, S., Krueger, F., Seconds-Pichon, A., Biggins, F., Wingett, S., 2015. FastQC. A quality control tool for high throughput sequence data. Babraham Bioinformatics. Babraham Inst.

Arnó, J., Castañé, C., Riudavets, J., Gabarra, R., 2010. Risk of damage to tomato crops by the generalist zoophytophagous predator Nesidiocoris tenuis (Reuter) (Hemiptera: Miridae). Bull. Entomol. Res. 100, 105–115. https://doi.org/10.1017/S0007485309006841

Banaei-Moghaddam, A.M., Meier, K., Karimi-Ashtiyani, R., Houben, A., 2013. Formation and expression of pseudogenes on the B chromosome of rye. Plant Cell 25, 2536–2544. https://doi.org/10.1105/tpc.113.111856

Benoit, J.B., Adelman, Z.N., Reinhardt, K., Dolan, A., Poelchau, M., Jennings, E.C., Szuter, E.M., Hagan, R.W., Gujar, H., Shukla, J.N., Zhu, F., Mohan, M., Nelson, D.R., Rosendale, A.J., Derst, C., Resnik, V., Wernig, S., Menegazzi, P., Wegener, C., Peschel, N., Hendershot, J.M., Blenau, W., Predel, R., Johnston, P.R., Ioannidis, P., Waterhouse, R.M., Nauen, R., Schorn, C., Ott, M.-C., Maiwald, F., Johnston, J.S., Gondhalekar, A.D., Scharf, M.E., Peterson, B.F., Raje, K.R., Hottel, B.A., Armisén, D., Crumière, A.J.J., Refki, P.N., Santos, M.E., Sghaier, E., Viala, S., Khila, A., Ahn, S.-J., Childers, C., Lee, C.-Y., Lin, H., Hughes, D.S.T., Duncan, E.J., Murali, S.C., Qu, J., Dugan, S., Lee, S.L., Chao, H., Dinh, H., Han, Y., Doddapaneni, H., Worley, K.C., Muzny, D.M., Wheeler, D., Panfilio, K.A., Vargas Jentzsch, I.M., Vargo, E.L., Booth, W., Friedrich, M., Weirauch, M.T., Anderson, M.A.E., Jones, J.W., Mittapalli, O., Zhao, C., Zhou, J.-J., Evans, J.D., Attardo, G.M., Robertson, H.M., Zdobnov, E.M., Ribeiro, J.M.C., Gibbs, R.A., Werren, J.H., Palli, S.R., Schal, C., Richards, S., 2016. Unique features of a global human ectoparasite identified through sequencing of the bed bug genome. Nat. Commun. 7, 10165. https://doi.org/10.1038/ncomms10165

Benson, G., 1999. Tandem repeats finder: A program to analyze DNA sequences. Nucleic Acids Res. 27, 573–580. https://doi.org/10.1093/nar/27.2.573

Biondi, A., Zappalà, L., Di Mauro, A., Tropea Garzia, G., Russo, A., Desneux, N., Siscaro, G., 2015. Can alternative host plant and prey affect phytophagy and biological control by the zoophytophagous mirid Nesidiocoris tenuis? BioControl 79–90. https://doi.org/10.1007/s10526-015-9700-5

Blankenfeldt, W., Kuzin, A.P., Skarina, T., Korniyenko, Y., Tong, L., Bayer, P., Janning, P., Thomashow, L.S., Mavrodi, D. V., 2004. Structure and function of the phenazine biosynthetic protein PhzF from Pseudomonas fluorescens. Proc. Natl. Acad. Sci. U. S. A. 101, 16431–16436. https://doi.org/10.1073/pnas.0407371101

Bolger, A.M., Lohse, M., Usadel, B., 2014. Trimmomatic: A flexible trimmer for Illumina sequence data. Bioinformatics 30, 2114–2120. https://doi.org/10.1093/bioinformatics/btu170

Bouagga, S., Urbaneja, A., Depalo, L., Rubio, L., Pérez-Hedo, M., 2019. Zoophytophagous predator-induced defences restrict accumulation of the tomato spotted wilt virus. Pest Manag. Sci. ps.5547. https://doi.org/10.1002/ps.5547

Boutet, E., Lieberherr, D., Tognolli, M., Schneider, M., Bairoch, A., 2008. UniProtKB/Swiss-Prot: The manually annotated section of the UniProt KnowledgeBase. Methods Mol. Biol. 406, 89–112. https://doi.org/10.1007/978-1-59745-535-0

Boyd, B.M., Allen, J.M., Koga, R., Fukatsu, T., Sweet, A.D., Johnson, K.P., Reed, D.L., 2016. Two Bacterial Genera, Sodalis and Rickettsia, Associated with the Seal Louse Proechinophthirus fluctus (Phthiraptera: Anoplura). Appl. Environ. Microbiol. 82, 3185– 3197. https://doi.org/10.1128/AEM.00282-16

Bruton, B.D., Mitchell, F., Fletcher, J., Pair, S.D., Wayadande, A., Melcher, U., Brady, J., Bextine, B., Popham, T.W., 2007. Serratia marcescens, a Phloem-Colonizing, Squash Bug - Transmitted Bacterium: Causal Agent of Cucurbit Yellow Vine Disease. Plant Dis. 87, 937– 944. https://doi.org/10.1094/pdis.2003.87.8.937

Calvo, J.F., Bolckmans, K., Stansly, P. a., Urbaneja, A., 2009. Predation by Nesidiocoris tenuis on Bemisia tabaci and injury to tomato. BioControl 54, 237–246. https://doi.org/10.1007/s10526-008-9164-y

Camacho, C., Coulouris, G., Avagyan, V., Ma, N., Papadopoulos, J., Bealer, K., Madden, T.L., 2009. BLAST+: Architecture and applications. BMC Bioinformatics 10, 421. https://doi.org/10.1186/1471-2105-10-421

Camacho, J.P.M., Sharbel, T.F., Beukeboom, L.W., 2000. B-chromosome evolution. Philos. Trans. R. Soc. London. Ser. B Biol. Sci. 355, 163–178. https://doi.org/10.1098/rstb.2000.0556

Carabajal Paladino, L.Z., Provazníková, I., Berger, M., Bass, C., Aratchige, N.S., López, S.N., Marec, F., Nguyen, P., 2019. Sex Chromosome Turnover in Moths of the Diverse Superfamily Gelechioidea. Genome Biol. Evol. 11, 1307–1319. https://doi.org/10.1093/gbe/evz075

Caspi-Fluger, A., Inbar, M., Steinberg, S., Friedmann, Y., Freund, M., Mozes-Daube, N., Zchori-Fein, E., 2014. Characterization of the symbiont Rickettsia in the mirid bug Nesidiocoris tenuis (Reuter) (Heteroptera: Miridae). Bull. Entomol. Res. 104, 681–688. https://doi.org/10.1017/S0007485314000492

Castañé, C., Arnó, J., Gabarra, R., Alomar, O., 2011. Plant damage to vegetable crops by zoophytophagous mirid predators. Biol. Control 59, 22–29. https://doi.org/10.1016/j.biocontrol.2011.03.007

Charlesworth, B., Charlesworth, D., 2000. The degeneration of Y chromosomes. Philos. Trans. R. Soc. B Biol. Sci. 355, 1563–1572. https://doi.org/10.1098/rstb.2000.0717

Chin, C.-S., Peluso, P., Sedlazeck, F.J., Nattestad, M., Concepcion, G.T., Clum, A., Dunn, C., O’Malley, R., Figueroa-Balderas, R., Morales-Cruz, A., Cramer, G.R., Delledonne, M., Luo, C., Ecker, J.R., Cantu, D., Rank, D.R., Schatz, M.C., 2016. Phased diploid genome assembly with single-molecule real-time sequencing. Nat. Methods 13, 1050–1054. https://doi.org/10.1038/nmeth.4035

Costechareyre, D., Balmand, S., Condemine, G., Rahbé, Y., 2012. Dickeya dadantii, a plant pathogenic bacterium producing cyt-like entomotoxins, causes septicemia in the pea aphid Acyrthosiphon pisum. PLoS One 7. https://doi.org/10.1371/journal.pone.0030702

Dai, X., Xun, H., Chang, J., Zhang, J., Hu, B., Li, H., Yuan, X., Cai, W., 2012. The complete mitochondrial genome of the plant bug Nesidiocoris tenuis (Reuter) (Hemiptera: Miridae: Bryocorinae: Dicyphini). Zootaxa 3554, 30–44. https://doi.org/10.11646/zootaxa.3554.1.2

Dalíková, M., Zrzavá, M., Hladová, I., Nguyen, P., Šonský, I., Flegrová, M., Kubíčková, S., Voleníková, A., Kawahara, A.Y., Peters, R.S., Marec, F., Sayres, M.W., 2017a. New Insights into the Evolution of the W Chromosome in Lepidoptera. J. Hered. 108, 709–719. https://doi.org/10.1093/jhered/esx063

Dalíková, M., Zrzavá, M., Kubíčková, S., Marec, F., 2017b. W-enriched satellite sequence in the Indian meal moth, Plodia interpunctella (Lepidoptera, Pyralidae). Chromosom. Res. 25, 241–252. https://doi.org/10.1007/s10577-017-9558-8

De Boer, J.G., Ode, P.J., Vet, L.E.M., Whitfield, J.B., Heimpel, G.E., 2007. Diploid males sire triploid daughters and sons in the parasitoid wasp Cotesia vestalis. Heredity (Edinb). 99, 288–294. https://doi.org/10.1038/sj.hdy.6800995

Doyle, JJ, Doyle, JL, 1990. Isolation of plant DNA from fresh tissue. Focus (Madison). 12, 13–15.

Dunning Hotopp, J.C., Clark, M.E., Oliveira, D.C.S.G., Foster, J.M., Fischer, P., Muñoz Torres, M.C., Giebel, J.D., Kumar, N., Ishmael, N., Wang, S., Ingram, J., Nene, R. V., Shepard, J., Tomkins, J., Richards, S., Spiro, D.J., Ghedin, E., Slatko, B.E., Tettelin, H., Werren, J.H., 2007. Widespread lateral gene transfer from intracellular bacteria to multicellular eukaryotes. Science (80-.). 317, 1753–1756. https://doi.org/10.1126/science.1142490

Ekblom, R., Wolf, J.B.W., 2014. A field guide to whole-genome sequencing, assembly and annotation. Evol. Appl. 1026–1042. https://doi.org/10.1111/eva.12178

Ellegren, H., 2014. Genome sequencing and population genomics in non-model organisms. Trends Ecol. Evol. 29, 51–63. https://doi.org/10.1016/j.tree.2013.09.008

Ferguson, K., 2020. Nesidiocoris tenuis PRJEB35378 linked-read genome annotation. https://doi.org/10.6084/m9.figshare.12073893.v1

Frydrychová, R., Grossmann, P., Trubac, P., Vítková, M., Marec, F., 2004. Phylogenetic distribution of TTAGG telomeric repeats in insects. Genome 47, 163–178. https://doi.org/10.1139/g03-100

Garantonakis, N., Pappas, M.L., Varikou, K., Skiada, V., Broufas, G.D., Kavroulakis, N., Papadopoulou, K.K., 2018. Tomato Inoculation With the Endophytic Strain Fusarium solani K Results in Reduced Feeding Damage by the Zoophytophagous Predator Nesidiocoris tenuis. Front. Ecol. Evol. 6, 1–7. https://doi.org/10.3389/fevo.2018.00126

Gatto, K.P., Mattos, J. V., Seger, K.R., Lourenço, L.B., 2018. Sex chromosome differentiation in the frog genus Pseudis involves satellite DNA and chromosome rearrangements. Front. Genet. 9, 1–12. https://doi.org/10.3389/fgene.2018.00301

Gomez-Rodriguez, V.M., Rodriguez-Garay, B., Palomino, G., Martínez, J., Barba-Gonzalez, R., 2013. Physical mapping of 5S and 18S ribosomal DNA in three species of Agave (Asparagales, Asparagaceae). Comp. Cytogenet. 7, 191–203. https://doi.org/10.3897/compcytogen.v7i3.5337

Grozeva, S., Anokhin, B.A., Simov, N., Kuznetsova, V.G., 2019. New evidence for the presence of the telomere motif (TTAGG) n in the family Reduviidae and its absence in the families Nabidae and Miridae (Hemiptera, Cimicomorpha). Comp. Cytogenet. 13, 283–295. https://doi.org/10.3897/compcytogen.v13i3.36676

Grozeva, S., Kuznetsova, V.G., Anokhin, B.A., 2011. Karyotypes, male meiosis and comparative FISH mapping of 18S ribosomal DNA and telomeric (TTAGG) n repeat in eight species of true bugs (Hemiptera, Heteroptera). Comp. Cytogenet. 5, 97–116. https://doi.org/10.3897/CompCytogen.v5i4.2307

Gurevich, A., Saveliev, V., Vyahhi, N., Tesler, G., 2013. QUAST: Quality assessment tool for genome assemblies. Bioinformatics 29, 1072–1075. https://doi.org/10.1093/bioinformatics/btt086

Haas, B.J., Salzberg, S.L., Zhu, W., Pertea, M., Allen, J.E., Orvis, J., White, O., Robin, C.R., Wortman, J.R., 2008. Automated eukaryotic gene structure annotation using EVidenceModeler and the Program to Assemble Spliced Alignments. Genome Biol. 9, 1– 22. https://doi.org/10.1186/gb-2008-9-1-r7

Hare, E.E., Johnston, J.S., 2011. Genome Size Determination Using Flow Cytometry of Propidium Iodide-Stained Nuclei, Molecular Methods for Evolutionary Genetics. https://doi.org/10.1007/978-1-61779-228-1

Huang, D.W., Sherman, B.T., Lempicki, R.A., 2009a. Bioinformatics enrichment tools: Paths toward the comprehensive functional analysis of large gene lists. Nucleic Acids Res. 37, 1–13. https://doi.org/10.1093/nar/gkn923

Huang, D.W., Sherman, B.T., Lempicki, R.A., 2009b. Systematic and integrative analysis of large gene lists using DAVID bioinformatics resources. Nat. Protoc. 4, 44–57. https://doi.org/10.1038/nprot.2008.211

Huang, H., McGarvey, P.B., Suzek, B.E., Mazumder, R., Zhang, J., Chen, Y., Wu, C.H., 2011. A comprehensive protein-centric ID mapping service for molecular data integration. Bioinformatics 27, 1190–1191. https://doi.org/10.1093/bioinformatics/btr101

Husnik, F., McCutcheon, J.P., 2018. Functional horizontal gene transfer from bacteria to eukaryotes. Nat. Rev. Microbiol. 16, 67–79. https://doi.org/10.1038/nrmicro.2017.137

Itou, M., Watanabe, M., Watanabe, E., Miura, K., 2013. Gut content analysis to study predatory efficacy of *Nesidiocoris tenuis* (Reuter) (Hemiptera: Miridae) by molecular methods. Entomol. Sci. 16, 145–150. https://doi.org/10.1111/j.1479-8298.2012.00552.x

Jang, E.B., Nishijima, K.A., 1990. Identification and Attractancy of Bacteria Associated with Dacus dorsalis (Diptera: Tephritidae). Environ. Entomol. 19, 1726–1731. https://doi.org/10.1093/ee/19.6.1726

Jauset, A.M., Edo-Tena, E., Castañé, C., Agustí, N., Alomar, O., Grozeva, S., 2015. Comparative cytogenetic study of three Macrolophus species (Heteroptera, Miridae). Comp. Cytogenet. 9, 613–623. https://doi.org/10.3897/CompCytogen.v9i4.5530

Jetybayev, I.Y., Bugrov, A.G., Ünal, M., Buleu, O.G., Rubtsov, N.B., 2017. Molecular cytogenetic analysis reveals the existence of two independent neo-XY sex chromosome systems in Anatolian Pamphagidae grasshoppers. BMC Evol. Biol. 17, 1–12. https://doi.org/10.1186/s12862-016-0868-9

Jones, R.N., Rees, H., 1982. B Chromosomes. Academic Press, New York.

Jones, Steven, Taylor, G., Chan, S., Warren, R., Hammond, S., Bilobram, S., Mordecai, G., Suttle, C., Miller, K., Schulze, A., Chan, A., Jones, Samantha, Tse, K., Li, I., Cheung, D., Mungall, K., Choo, C., Ally, A., Dhalla, N., Tam, A., Troussard, A., Kirk, H., Pandoh, P., Paulino, D., Coope, R., Mungall, A., Moore, R., Zhao, Y., Birol, I., Ma, Y., Marra, M., Haulena, M., 2017. The Genome of the Beluga Whale (Delphinapterus leucas). Genes (Basel). 8, 378. https://doi.org/10.3390/genes8120378

Jung, S., Lee, S., 2012. Molecular phylogeny of the plant bugs (Heteroptera: Miridae) and the evolution of feeding habits. Cladistics 28, 50–79. https://doi.org/10.1111/j.1096-0031.2011.00365.x

Kamber, T., Smits, T.H.M., Rezzonico, F., Duffy, B., 2012. Genomics and current genetic understanding of Erwinia amylovora and the fire blight antagonist Pantoea vagans. Trees - Struct. Funct. 26, 227–238. https://doi.org/10.1007/s00468-011-0619-x

Kato, A., Albert, P.S., Vega, J.M., Birchler, J.A., 2006. Sensitive fluorescence in situ hybridization signal detection in maize using directly labeled probes produced by high concentration DNA polymerase nick translation. Biotech. Histochem. 81, 71–78. https://doi.org/10.1080/10520290600643677

Keeling, C.I., Yuen, M.M.S., Liao, N.Y., Roderick Docking, T., Chan, S.K., Taylor, G.A., Palmquist, D.L., Jackman, S.D., Nguyen, A., Li, M., Henderson, H., Janes, J.K., Zhao, Y., Pandoh, P., Moore, R., Sperling, F.A.H., W Huber, D.P., Birol, I., Jones, S.J.M., Bohlmann, J., 2013. Draft genome of the mountain pine beetle, Dendroctonus ponderosae Hopkins, a major forest pest. Genome Biol. 14, R27. https://doi.org/10.1186/gb-2013-14-3-r27

Keilwagen, J., Wenk, M., Erickson, J.L., Schattat, M.H., Grau, J., Hartung, F., 2016. Using intron position conservation for homology-based gene prediction. Nucleic Acids Res. 44. https://doi.org/10.1093/nar/gkw092

Kofler, R., Orozco-terWengel, P., de Maio, N., Pandey, R.V., Nolte, V., Futschik, A., Kosiol, C., Schlötterer, C., 2011. Popoolation: A toolbox for population genetic analysis of next generation sequencing data from pooled individuals. PLoS One 6. https://doi.org/10.1371/journal.pone.0015925

Kuznetsova, V.G., Grozeva, S.M., Nokkala, S., Nokkala, C., 2011. Cytogenetics of the true bug infraorder cimicomorpha (hemiptera, heteroptera): A review. Zookeys 154, 31–70. https://doi.org/10.3897/zookeys.154.1953

Lee, W., Kang, J., Jung, C., Hoelmer, K., Lee, S.H., Lee, S., 2009. Complete mitochondrial genome of brown marmorated stink bug Halyomorpha halys (Hemiptera: Pentatomidae), and phylogenetic relationships of hemipteran suborders. Mol. Cells 28, 155–165. https://doi.org/10.1007/s10059-009-0125-9

Legeai, F., Shigenobu, S., Gauthier, J.P., Colbourne, J., Rispe, C., Collin, O., Richards, S., Wilson, A.C.C., Murphy, T., Tagu, D., 2010. AphidBase: A centralized bioinformatic resource for annotation of the pea aphid genome. Insect Mol. Biol. 19, 5–12. https://doi.org/10.1111/j.1365-2583.2009.00930.x

Leung, K., Ras, E., Ferguson, K.B., Ariëns, S., Babendreier, D.B., Bijma, P., Bourtzis, K., Brodeur, J., Bruins, M., Centurión, A., Chattington, S., Chinchilla-Ramírez, M., Dicke, M., Fatouros, N., González Cabrera, J., Groot, T., Haye, T., Knapp, M., Koskinioti, P., Le Hesran, S., Lirakis, M., Paspati, A., Pérez-Hedo, M., Plouvier, W., Schlötterer, C., Stahl, J., Thiel, A., Urbaneja, A., van de Zande, L., Verhulst, E., Vet, L., Visser, S., Werren, J., Xia, S., Zwaan, B., Magalhães, S., Beukeboom, L., Pannebakker, B., 2019. Next Generation Biological Control: the Need for Integrating Genetics and Evolution. Preprints 1–34. https://doi.org/10.20944/preprints201911.0300.v1

Li, H., Handsaker, B., Wysoker, A., Fennell, T., Ruan, J., Homer, N., Marth, G., Abecasis, G., Durbin, R., 2009. The Sequence Alignment/Map format and SAMtools. Bioinformatics 25, 2078–2079. https://doi.org/10.1093/bioinformatics/btp352

Lindsey, A.R.I., Kelkar, Y.D., Wu, X., Sun, D., Martinson, E.O., Yan, Z., Rugman-Jones, P.F., Hughes, D.S.T., Murali, S.C., Qu, J., Dugan, S., Lee, S.L., Chao, H., Dinh, H., Han, Y., Doddapaneni, H.V., Worley, K.C., Muzny, D.M., Ye, G., Gibbs, R.A., Richards, S., Yi, S. V., Stouthamer, R., Werren, J.H., 2018. Comparative genomics of the miniature wasp and pest control agent Trichogramma pretiosum. BMC Biol. 16, 1–20. https://doi.org/10.1186/s12915-018-0520-9

Lindsey, A.R.I., Werren, J.H., Richards, S., Stouthamer, R., 2016. Comparative Genomics of a Parthenogenesis-Inducing Wolbachia Symbiont. G3 Genes, Genomes, Genet. 6, 2113–2123. https://doi.org/10.1534/g3.116.028449

Lowe-Power, T.M., Khokhani, D., Allen, C., 2018. How Ralstonia solanacearum Exploits and Thrives in the Flowing Plant Xylem Environment. Trends Microbiol. 26, 929–942. https://doi.org/10.1016/j.tim.2018.06.002

Majoros, W.H., Pertea, M., Salzberg, S.L., 2004. TigrScan and GlimmerHMM: Two open source ab initio eukaryotic gene-finders. Bioinformatics 20, 2878–2879. https://doi.org/10.1093/bioinformatics/bth315

Marçais, G., Kingsford, C., 2011. A fast, lock-free approach for efficient parallel counting of occurrences of k-mers. Bioinformatics 27, 764–770. https://doi.org/10.1093/bioinformatics/btr011

Marec, F., Shvedov, A.N., 1990. Yellow eye, a new pigment mutation in Ephestia kuehniella Zeller (Lepidoptera: Pyralidae). Hereditas 113, 97–100. https://doi.org/10.1111/j.1601-5223.1990.tb00072.x

Martínez-García, H., Román-Fernández, L.R., Sáenz-Romo, M.G., Pérez-Moreno, I., Marco-Mancebón, V.S., 2016. Optimizing Nesidiocoris tenuis (Hemiptera: Miridae) as a biological control agent: mathematical models for predicting its development as a function of temperature. Bull. Entomol. Res. 106, 215–224. https://doi.org/10.1017/S0007485315000978

May, C.M., Heuvel, J., Doroszuk, A., Hoedjes, K.M., Flatt, T., Zwaan, B.J., 2019. Adaptation to developmental diet influences the response to selection on age at reproduction in the fruit fly. J. Evol. Biol. 32, 425–437. https://doi.org/10.1111/jeb.13425

Mediouni, J., Fuková, I., Frydrychová, R., Marec, F., Fuková, I., Marec, F., Dhouibi, M.H., 2004. Karyotype, sex chromatin and sex chromosome differentiation in the carob moth, Ectomyelois ceratoniae (Lepidoptera: Pyralidae). Caryologia 57, 184–194. https://doi.org/10.1080/00087114.2004.10589391

Mollá, O., Biondi, A., Alonso-Valiente, M., Urbaneja, A., 2014. A comparative life history study of two mirid bugs preying on Tuta absoluta and Ephestia kuehniella eggs on tomato crops: Implications for biological control. BioControl 59, 175–183. https://doi.org/10.1007/s10526-013-9553-8

Morgulis, A., Gertz, E.M., Schäffer, A.A., Agarwala, R., 2006. A fast and symmetric DUST implementation to mask low-complexity DNA sequences. J. Comput. Biol. 13, 1028–1040. https://doi.org/10.1089/cmb.2006.13.1028

Novák, P., Neumann, P., Pech, J., Steinhaisl, J., MacAs, J., 2013. RepeatExplorer: A Galaxy-based web server for genome-wide characterization of eukaryotic repetitive elements from next-generation sequence reads. Bioinformatics 29, 792–793. https://doi.org/10.1093/bioinformatics/btt054

Panfilio, K.A., Angelini, D.R., 2018. By land, air, and sea: hemipteran diversity through the genomic lens. Curr. Opin. Insect Sci. 25, 106–115. https://doi.org/10.1016/j.cois.2017.12.005

Panfilio, K.A., Vargas Jentzsch, I.M., Benoit, J.B., Erezyilmaz, D., Suzuki, Y., Colella, S., Robertson, H.M., Poelchau, M.F., Waterhouse, R.M., Ioannidis, P., Weirauch, M.T., Hughes, D.S.T., Murali, S.C., Werren, J.H., Jacobs, C.G.C., Duncan, E.J., Armisén, D., Vreede, B.M.I., Baa-Puyoulet, P., Berger, C.S., Chang, C., Chao, H., Chen, M.-J.M., Chen, Y.-T., Childers, C.P., Chipman, A.D., Cridge, A.G., Crumière, A.J.J., Dearden, P.K., Didion, E.M., Dinh, H., Doddapaneni, H.V., Dolan, A., Dugan, S., Extavour, C.G., Febvay, G., Friedrich, M., Ginzburg, N., Han, Y., Heger, P., Holmes, C.J., Horn, T., Hsiao, Y., Jennings, E.C., Johnston, J.S., Jones, T.E., Jones, J.W., Khila, A., Koelzer, S., Kovacova, V., Leask, M., Lee, S.L., Lee, C.-Y., Lovegrove, M.R., Lu, H., Lu, Y., Moore, P.J., Munoz-Torres, M.C., Muzny, D.M., Palli, S.R., Parisot, N., Pick, L., Porter, M.L., Qu, J., Refki, P.N., Richter, R., Rivera-Pomar, R., Rosendale, A.J., Roth, S., Sachs, L., Santos, M.E., Seibert, J., Sghaier, E., Shukla, J.N., Stancliffe, R.J., Tidswell, O., Traverso, L., van der Zee, M., Viala, S., Worley, K.C., Zdobnov, E.M., Gibbs, R.A., Richards, S., 2019. Molecular evolutionary trends and feeding ecology diversification in the Hemiptera, anchored by the milkweed bug genome. Genome Biol. 20, 64. https://doi.org/10.1186/s13059-019-1660-0

Paspati, A., Ferguson, K.B., Verhulst, E.C., Urbaneja, A., González-Cabrera, J., Pannebakker, B.A., 2019. Effect of mass rearing on the genetic diversity of the predatory mite Amblyseius swirskii Athias-Henriot (Acari: Phytoseiidae). Entomol. Exp. Appl. 167, 670–681. https://doi.org/10.1111/eea.12811

Pérez-Hedo, M., Arias-Sanguino, Á.M., Urbaneja, A., 2018. Induced tomato plant resistance against tetranychus urticae triggered by the phytophagy of nesidiocoris tenuis. Front. Plant Sci. 9, 1–8. https://doi.org/10.3389/fpls.2018.01419

Pérez-Hedo, M., Urbaneja-Bernat, P., Jaques, J.A., Flors, V., Urbaneja, A., 2015. Defensive plant responses induced by Nesidiocoris tenuis (Hemiptera: Miridae) on tomato plants. J. Pest Sci. (2004). 88, 543–554. https://doi.org/10.1007/s10340-014-0640-0

Pérez-Hedo, M., Urbaneja, A., 2016. The Zoophytophagous Predator Nesidiocoris tenuis: A Successful But Controversial Biocontrol Agent in Tomato Crops, in: Advances in Insect Control and Resistance Management. Springer International Publishing, Cham, pp. 121– 138. https://doi.org/10.1007/978-3-319-31800-4_7

Pita, S., Panzera, F., Mora, P., Vela, J., Palomeque, T., Lorite, P., 2016. The presence of the ancestral insect telomeric motif in kissing bugs (Triatominae) rules out the hypothesis of its loss in evolutionarily advanced Heteroptera (Cimicomorpha). Comp. Cytogenet. 10, 427–437. https://doi.org/10.3897/compcytogen.v10i3.9960

Poelchau, M., Childers, C., Moore, G., Tsavatapalli, V., Evans, J., Lee, C.Y., Lin, H., Lin, J.W., Hackett, K., 2015. The i5k Workspace@NAL-enabling genomic data access, visualization and curation of arthropod genomes. Nucleic Acids Res. 43, D714–D719. https://doi.org/10.1093/nar/gku983

Prado, S.S., Almeida, R.P.P., 2009. Phylogenetic placement of pentatomid stink bug gut symbionts. Curr. Microbiol. 58, 64–69. https://doi.org/10.1007/s00284-008-9267-9

Puentes, A., Stephan, J.G., Björkman, C., 2018. A Systematic Review on the Effects of Plant-Feeding by Omnivorous Arthropods: Time to Catch-Up With the Mirid-Tomato Bias? Front. Ecol. Evol. 6. https://doi.org/10.3389/fevo.2018.00218

Quan, Q., Hu, X., Pan, B., Zeng, B., Wu, N., Fang, G., Cao, Y., Chen, X., Li, X., Huang, Y., Zhan, S., 2019. Draft genome of the cotton aphid Aphis gossypii. Insect Biochem. Mol. Biol. 105, 25–32. https://doi.org/10.1016/j.ibmb.2018.12.007

Rasmussen, L.B., Jensen, K., Sørensen, J.G., Sverrisdóttir, E., Nielsen, K.L., Overgaard, J., Holmstrup, M., Kristensen, T.N., 2018. Are commercial stocks of biological control agents genetically depauperate? – A case study on the pirate bug Orius majusculus Reuter. Biol. Control 127, 31–38. https://doi.org/10.1016/j.biocontrol.2018.08.016

Reddiex, A.J., Allen, S.L., Chenoweth, S.F., 2018. A Genomic Reference Panel for Drosophila serrata. G3 Genes, Genomes, Genet. 8, 1335–1346. https://doi.org/10.1534/g3.117.300487

Richards, S., Gibbs, R.A., Gerardo, N.M., Moran, N., Nakabachi, A., Stern, D., Tagu, D., Wilson, A.C.C., Muzny, D., Kovar, C., Cree, A., Chacko, J., Chandrabose, M.N., Dao, M.D., Dinh, H.H., Gabisi, R.A., Hines, S., Hume, J., Jhangian, S.N., Joshi, V., Lewis, L.R., Liu, Y.S., Lopez, J., Morgan, M.B., Nguyen, N.B., Okwuonu, G.O., Ruiz, S.J., Santibanez, J., Wright, R.A., Fowler, G.R., Hitchens, M.E., Lozado, R.J., Moen, C., Steffen, D., Warren, J.T., Zhang, J., Nazareth, L. V., Chavez, D., Davis, C., Lee, S.L., Patel, B.M., Pu, L.L., Bell, S.N., Johnson, A.J., Vattathil, S., Williams, R.L., Shigenobu, S., Dang, P.M., Morioka, M., Fukatsu, T., Kudo, T., Miyagishima, S.Y., Jiang, H., Worley, K.C., Legeai, F., Gauthier, J.P., Collin, O., Zhang, L., Chen, H.C., Ermolaeva, O., Hlavina, W., Kapustin, Y., Kiryutin, B., Kitts, P., Maglott, D., Murphy, T., Pruitt, K., Sapojnikov, V., Souvorov, A., Thibaud-Nissen, F., Câmara, F., Guigó, R., Stanke, M., Solovyev, V., Kosarev, P., Gilbert, D., Gabaldón, T., Huerta-Cepas, J., Marcet-Houben, M., Pignatelli, M., Moya, A., Rispe, C., Ollivier, M., Quesneville, H., Permal, E., Llorens, C., Futami, R., Hedges, D., Robertson, H.M., Alioto, T., Mariotti, M., Nikoh, N., McCutcheon, J.P., Burke, G., Kamins, A., Latorre, A., Ashton, P., Calevro, F., Charles, H., Colella, S., Douglas, A.E., Jander, G., Jones, D.H., Febvay, G., Kamphuis, L.G., Kushlan, P.F., Macdonald, S., Ramsey, J., Schwartz, J., Seah, S., Thomas, G., Vellozo, A., Cass, B., Degnan, P., Hurwitz, B., Leonardo, T., Koga, R., Altincicek, B., Anselme, C., Atamian, H., Barribeau, S.M., De Vos, M., Duncan, E.J., Evans, J., Ghanim, M., Heddi, A., Kaloshian, I., Vincent-Monegat, C., Parker, B.J., Pérez-Brocal, V., Rahbé, Y., Spragg, C.J., Tamames, J., Tamarit, D., Tamborindeguy, C., Vilcinskas, A., Bickel, R.D., Brisson, J.A., Butts, T., Chang, C.C., Christiaens, O., Davis, G.K., Duncan, E., Ferrier, D., Iga, M., Janssen, R., Lu, H.L., McGregor, A., Miura, T., Smagghe, G., Smith, J., Van Der Zee, M., Velarde, R., Wilson, M., Dearden, P., Edwards, O.R., Gordon, K., Hilgarth, R.S., Rider, S.D., Srinivasan, D., Walsh, T.K., Ishikawa, A., Jaubert-Possamai, S., Fenton, B., Huang, W., Rizk, G., Lavenier, D., Nicolas, J., Smadja, C., Zhou, J.J., Vieira, F.G., He, X.L., Liu, R., Rozas, J., Field, L.M., Campbell, P., Carolan, J.C., Fitzroy, C.I.J., Reardon, K.T., Reeck, G.R., Singh, K., Wilkinson, T.L., Huybrechts, J., Abdel-Latief, M., Robichon, A., Veenstra, J.A., Hauser, F., Cazzamali, G., Schneider, M., Williamson, M., Stafflinger, E., Hansen, K.K., Grimmelikhuijzen, C.J.P., Price, D.R.G., Caillaud, M., Van Fleet, E., Ren, Q., Gatehouse, J.A., Brault, V., Monsion, B., Diaz, J., Hunnicutt, L., Ju, H.J., Pechuan, X., Aguilar, J., Cortés, T., Ortiz-Rivas, B., Martínez-Torres, D., Dombrovsky, A., Dale, R.P., Davies, T.G.E., Williamson, M.S., Jones, A., Sattelle, D., Williamson, S., Wolstenholme, A., Cottret, L., Sagot, M.F., Heckel, D.G., Hunter, W., 2010. Genome sequence of the pea aphid Acyrthosiphon pisum. PLoS Biol. 8. https://doi.org/10.1371/journal.pbio.1000313

Richards, S., Murali, S.C., 2015. Best practices in insect genome sequencing: what works and what doesn’t. Curr. Opin. Insect Sci. 7, 1–7. https://doi.org/10.1016/j.cois.2015.02.013

Sachman-Ruiz, B., Quiroz-Castañeda, R.E., 2018. Genomics of Rickettsiaceae: An Update, in: Farm Animals Diseases, Recent Omic Trends and New Strategies of Treatment. InTech, p. 13. https://doi.org/10.5772/intechopen.74563

Sahara, K., Marec, F., Traut, W., 1999. TTAGG telomeric repeats in chromosomes of some insects and other arthropods. Chromosom. Res. 7, 449–460. https://doi.org/10.1023/A:1009297729547

Sanchez, J.A., 2009. Density thresholds for Nesidiocoris tenuis (Heteroptera: Miridae) in tomato crops. Biol. Control 51, 493–498. https://doi.org/10.1016/j.biocontrol.2009.09.006

Santos-Garcia, D., Silva, F.J., Morin, S., Dettner, K., Kuechler, S.M., 2017. The all-rounder sodalis: A new bacteriome-associated endosymbiont of the lygaeoid bug henestaris halophilus (heteroptera: Henestarinae) and a critical examination of its evolution. Genome Biol. Evol. 9, 2893–2910. https://doi.org/10.1093/gbe/evx202

Schwarz, A., Medrano-Mercado, N., Schaub, G.A., Struchiner, C.J., Bargues, M.D., Levy, M.Z., Ribeiro, J.M.C., 2014. An Updated Insight into the Sialotranscriptome of Triatoma infestans: Developmental Stage and Geographic Variations. PLoS Negl. Trop. Dis. 8. https://doi.org/10.1371/journal.pntd.0003372

Sember, A., Bertollo, L.A.C., Ráb, P., Yano, C.F., Hatanaka, T., de Oliveira, E.A., Cioffi, M. de B., 2018. Sex chromosome evolution and genomic divergence in the fish Hoplias malabaricus (Characiformes, Erythrinidae). Front. Genet. 9, 1–12. https://doi.org/10.3389/fgene.2018.00071

Simão, F.A., Waterhouse, R.M., Ioannidis, P., Kriventseva, E. V., Zdobnov, E.M., 2015. BUSCO: Assessing genome assembly and annotation completeness with single-copy orthologs. Bioinformatics 31, 3210–3212. https://doi.org/10.1093/bioinformatics/btv351

Sobreira, T.J.P., Durham, A.M., Gruber, A., 2006. TRAP: Automated classification, quantification and annotation of tandemly repeated sequences. Bioinformatics 22, 361–362. https://doi.org/10.1093/bioinformatics/bti809

Sochorová, J., Garcia, S., Gálvez, F., Symonová, R., Kovařík, A., 2018. Evolutionary trends in animal ribosomal DNA loci: introduction to a new online database. Chromosoma 127, 141–150. https://doi.org/10.1007/s00412-017-0651-8

Streito, J.C., Clouet, C., Hamdi, F., Gauthier, N., 2017. Population genetic structure of the biological control agent Macrolophus pygmaeus in Mediterranean agroecosystems. Insect Sci. 24, 859–876. https://doi.org/10.1111/1744-7917.12370

Sunnucks, P., Hales, D.F., 1996. Numerous transposed sequences of mitochondrial cytochrome oxidase I-II in aphids of the genus Sitobion (Hemiptera: Aphididae). Mol. Biol. Evol. 13, 510–524. https://doi.org/10.1093/oxfordjournals.molbev.a025612

Szűcs, M., Vercken, E., Bitume, E. V., Hufbauer, R.A., 2019. The implications of rapid eco-evolutionary processes for biological control - a review. Entomol. Exp. Appl. eea.12807. https://doi.org/10.1111/eea.12807

Testa, A.C., Hane, J.K., Ellwood, S.R., Oliver, R.P., 2015. CodingQuarry: Highly accurate hidden Markov model gene prediction in fungal genomes using RNA-seq transcripts. BMC Genomics 16, 1–12. https://doi.org/10.1186/s12864-015-1344-4

Thomas, G.W.C., Dohmen, E., Hughes, D.S.T., Murali, S.C., Poelchau, M., Glastad, K., Anstead, C.A., Ayoub, N.A., Batterham, P., Bellair, M., Binford, G.J., Chao, H., Chen, Y.H., Childers, C., Dinh, H., Doddapaneni, H.V., Duan, J.J., Dugan, S., Esposito, L.A., Friedrich, M., Garb, J., Gasser, R.B., Goodisman, M.A.D., Gundersen-Rindal, D.E., Han, Y., Handler, A.M., Hatakeyama, M., Hering, L., Hunter, W.B., Ioannidis, P., Jayaseelan, J.C., Kalra, D., Khila, A., Korhonen, P.K., Lee, C.E., Lee, S.L., Li, Y., Lindsey, A.R.I., Mayer, G., McGregor, A.P., McKenna, D.D., Misof, B., Munidasa, M., Munoz-Torres, M., Muzny, D.M., Niehuis, O., Osuji-Lacy, N., Palli, S.R., Panfilio, K.A., Pechmann, M., Perry, T., Peters, R.S., Poynton, H.C., Prpic, N.-M., Qu, J., Rotenberg, D., Schal, C., Schoville, S.D., Scully, E.D., Skinner, E., Sloan, D.B., Stouthamer, R., Strand, M.R., Szucsich, N.U., Wijeratne, A., Young, N.D., Zattara, E.E., Benoit, J.B., Zdobnov, E.M., Pfrender, M.E., Hackett, K.J., Werren, J.H., Worley, K.C., Gibbs, R.A., Chipman, A.D., Waterhouse, R.M., Bornberg-Bauer, E., Hahn, M.W., Richards, S., 2020. Gene content evolution in the arthropods. Genome Biol. 21, 15. https://doi.org/10.1186/s13059-019-1925-7

Tian, C., Tek Tay, W., Feng, H., Wang, Y., Hu, Y., Li, G., 2015. Characterization of Adelphocoris suturalis (Hemiptera: Miridae) Transcriptome from Different Developmental Stages. Sci. Rep. 5, 11042. https://doi.org/10.1038/srep11042

Trapnell, C., Pachter, L., Salzberg, S.L., 2009. TopHat: Discovering splice junctions with RNA-Seq. Bioinformatics 25, 1105–1111. https://doi.org/10.1093/bioinformatics/btp120

Trapnell, C., Williams, B.A., Pertea, G., Mortazavi, A., Kwan, G., van Baren, M.J., Salzberg, S.L., Wold, B.J., Pachter, L., 2010. Transcript assembly and quantification by RNA-Seq reveals unannotated transcripts and isoform switching during cell differentiation. Nat. Biotechnol. 28, 511–5. https://doi.org/10.1038/nbt.1621

Traut, W., 1976. Pachytene mapping in the female silkworm, Bombyx mori L. (Lepidoptera). Chromosoma 58, 275–284. https://doi.org/10.1007/BF00292094

Traut, W., Sahara, K., Otto, T.D., Marec, F., 1999. Molecular differentiation of sex chromosomes probed by comparative genomic hybridization. Chromosoma 108, 173–180. https://doi.org/10.1007/s004120050366

Traverse, K.L., Pardue, M.L., 1988. A spontaneously opened ring chromosome of Drosophila melanogaster has acquired He-T DNA sequences at both new telomeres. Proc. Natl. Acad. Sci. U. S. A. 85, 8116–8120. https://doi.org/10.1073/pnas.85.21.8116

Urbaneja-Bernat, P., Bru, P., González-Cabrera, J., Urbaneja, A., Tena, A., 2019. Reduced phytophagy in sugar-provisioned mirids. J. Pest Sci. (2004). 92, 1139–1148. https://doi.org/10.1007/s10340-019-01105-9

Urbaneja, A., González-Cabrera, J., Arnó, J., Gabarra, R., 2012. Prospects for the biological control of Tuta absoluta in tomatoes of the Mediterranean basin. Pest Manag. Sci. 68, 1215–1222. https://doi.org/10.1002/ps.3344

van Lenteren, J.C., Bolckmans, K., Köhl, J., Ravensberg, W.J., Urbaneja, A., 2018. Biological control using invertebrates and microorganisms: plenty of new opportunities. BioControl 63, 39–59. https://doi.org/10.1007/s10526-017-9801-4

van Wilgenburg, E., Driessen, G., Beukeboom, L.W., 2006. Single locus complementary sex determination in Hymenoptera: an “unintelligent” design? Front. Zool. 3, 1. https://doi.org/https://doi.org/10.1186/1742-9994-3-1

Vurture, G.W., Sedlazeck, F.J., Nattestad, M., Schatz, M.C., Gurtowski, J., Underwood, C.J., Vurture, G.W., Fang, H., 2017. GenomeScope: fast reference-free genome profiling from short reads. Bioinformatics 33, 2202–2204. https://doi.org/10.1093/bioinformatics/btx153

Weisenfeld, N.I., Kumar, V., Shah, P., Church, D.M., Jaffe, D.B., 2017. Direct determination of diploid genome sequences. Genome Res. 27, 757–767. https://doi.org/10.1101/gr.214874.116.Freely

Wheeler, D., Redding, A.J., Werren, J.H., 2013. Characterization of an Ancient Lepidopteran Lateral Gene Transfer. PLoS One 8. https://doi.org/10.1371/journal.pone.0059262

Xu, L., Dong, Z., Fang, L., Luo, Y., Wei, Z., Guo, H., Zhang, G., Gu, Y.Q., Coleman-Derr, D., Xia, Q., Wang, Y., 2019. OrthoVenn2: A web server for genome wide comparison and annotation of orthologous clusters across multiple species. Nucleic Acids Res. 47, W52– W58. https://doi.org/10.1093/nar/gkz333

Xu, P., Lu, B., Liu, J., Chao, J., Donkersley, P., Holdbrook, R., Lu, Y., 2019. Duplication and expression of horizontally transferred polygalacturonase genes is associated with host range expansion of mirid bugs. BMC Evol. Biol. 19, 12. https://doi.org/10.1186/s12862-019-1351-1

Xun, H., Li, H., Li, S., Wei, S., Zhang, L., Song, F., Jiang, P., Yang, H., Han, F., Cai, W., 2016. Population genetic structure and post-LGM expansion of the plant bug Nesidiocoris tenuis (Hemiptera: Miridae) in China. Sci. Rep. 6, 26755. https://doi.org/10.1038/srep26755

Zhang, Y., Qiu, S., 2015. Examining phylogenetic relationships of Erwinia and Pantoea species using whole genome sequence data. Antonie van Leeuwenhoek, Int. J. Gen. Mol. Microbiol. 108, 1037–1046. https://doi.org/10.1007/s10482-015-0556-6

Zhou, W., Rousset, F., O’Neill, S., 1998. Phylogeny and PCR-based classification of Wolbachia strains using wsp gene sequences. Proc. R. Soc. B Biol. Sci. 265, 509–515. https://doi.org/10.1098/rspb.1998.0324

Zrzavá, M., Hladová, I., Dalíková, M., Šíchová, J., Õunap, E., Kubíčková, S., Marec, F., 2018. Sex chromosomes of the iconic moth Abraxas grossulariata (Lepidoptera, Geometridae) and its congener A. sylvata. Genes (Basel). 9, 1–16. https://doi.org/10.3390/genes9060279

