## supplementary materials for "Jekyll or Hyde? The genome (and more) of *Nesidiocoris tenuis*, a zoophytophagous predatory bug that is both a biological control agent and a pest"

*Nesidiocoris tenuis* genome – Supplementary information S1

For DANS EASY Repository data:

Pannebakker, dr. ir. B. A. (Wageningen University); Ferguson, K. B. (Wageningen University) (2019): The *Nesidiocoris tenuis* genome manuscript supporting data. DANS. https://doi.org/10.17026/dans-z5z-zec9

#### S1.1. *Wolbachia* PCR results

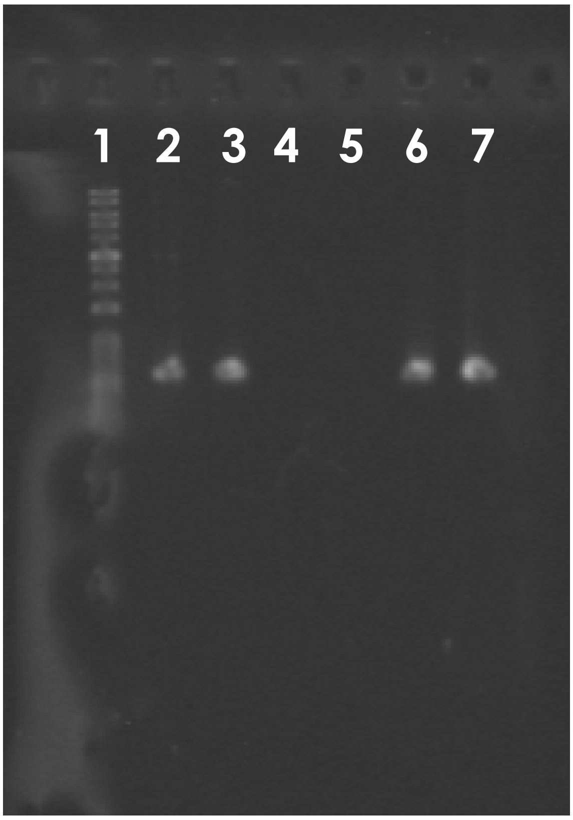

##### Figure S1.1.1

*Wolbachia* detection was carried out according to the PCR protocol of Zhou et al. (Zhou, Rousset, and Neill 1998). Lane 1 contains a 100bp ladder, while lanes 2 and 3 contain PCR products for *N. tenuis* DNA (see 18S procedure). Lanes 4 and 5 contain sterilized water as a negative control, while lanes 6 and 7 contain PCR products for *E. kuehniella* DNA (see Southern Blot procedure).

#### S1.2. Flow cytometry data

Files associated with the flow cytometry data collected to determine genome size. The .FCS files of the results include: two PI-stained *N. tenuis* samples, one sample of PI-stained *N. tenuis* dosed with head of *D. melanogaster*, one unstained *N. tenuis* background sample, and three PI-stained *D. melanogaster* samples. Available on DANS in collection, refer to these files:

- original/20160226_26_Nesidiocoristenuis1.fcs
- original/20160226_27_Nesidiocoristenuis2.fcs
- original/20160226_28_NesidiocoristenuisDMelanogaster.fcs
- original/20160226_11_unstained_Nesidiocoristenuis.fcs
- original/20160226_Dmelanogaster_PI stained_3.fcs
- original/20160226_DMelanogaster_PI stained_4.fcs
- original/20160226_DMelanogaster_PI stained_5.fcs

#### S1.3. Decontamination and potential LGT identification

S1.3.1 In-house reference bacterial database that contains 2,100 different bacterial species. Adapted from (Wheeler, Redding, and Werren 2013). Available on DANS:

- original/S1.3.1_LGTPipeline_BacteriaSet.csv

S1.3.2 List of bacterial contamination, identified an adapted pipeline (Wheeler, Redding, and Werren 2013). Scaffold numbers correspond to removed scaffolds in S1.3.3. Available on DANS:

- original/S1.3.2_Nesidiocoris_BacterialScaffolds.txt

S1.3.3. The .FASTA files of scaffolds deemed contamination. Note, the names are no longer valid when compared to the current assembly. Available on DANS:

- original/S1.3.3_Contamination.fa

S1.3.4 List of potential LGT candidates, limited selection, available on DANS:

- original/S1.3.4_Nesidiocoris_LGTsManual_Limited.csv

#### S1.4. Gene List (UniProtKB list) and DAVID Reports

A table displaying the gene IDs from accession numbers in the assembly (excepting "No_blast_hit" or those that did not map to a UniProtKB ID), the final UniProtKB Accession IDs, and the deduplicated list that was used with DAVID, available on DANS:

- original/AnnotationToUniProtKBtoDAVID.ods

The full table of the larger DAVID Gene List Report and the smaller DAVID Gene Report is available on DANS:

- original/DAVIDGeneListReport.txt
- original/DAVIDGeneReport.txt

#### S1.5. Full protein set

The .faa (amino acid .fasta) used in the protein cluster analysis is the official protein set of the *N. tenuis* genome and is available on DANS.

- original/Ntenuis_protein.aa.fasta

#### S1.6. Cytogenetics: Southern blot, abundance distributions, and primers

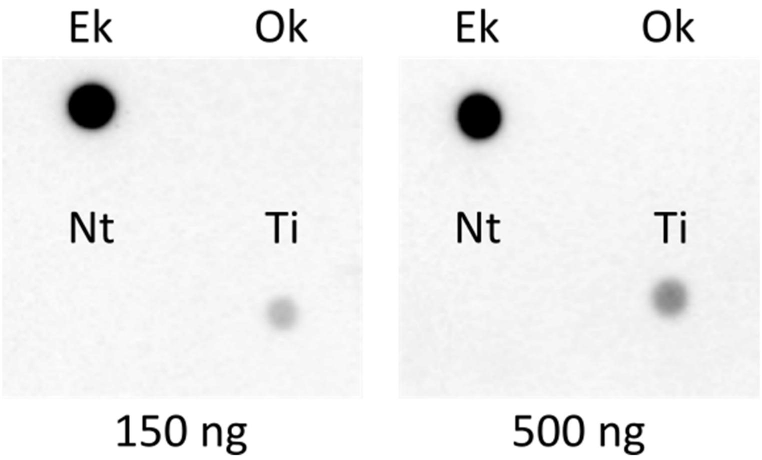

##### Figure S1.6.1.

Southern dot blot assay using the insect telomere motif $(TTAGG)_n$ in *E. kuehniella*, *O. keta*, *N. tenuis*, and *T. infestans*. *E. kuehniella* (Ek) and *T. infestans* (Ti) show hybridization signals, whilst *N. tenuis* (Nt) and the negative control *O. keta* (Ok) display no detectable hybridization signals. Both quantities of DNA (150 ng and 500 ng) show comparable results.

Table S1.6.1. Abundance and distribution of Nt_rep1 in the assembled genome

| **# of Nt_rep1 copies per scaffold** | **Number of scaffolds in the assembly** | **% of total** |
|---|---|---|
| 1 | 2625 | 78.45 |
| 2 | 441 | 13.18 |
| 3 | 148 | 4.42 |
| 4 | 74 | 2.21 |
| 5 | 32 | 0.96 |
| 6 | 11 | 0.33 |
| 7 | 3 | 0.09 |
| 8 | 6 | 0.18 |
| 9 | 3 | 0.09 |
| 11 | 2 | 0.06 |
| 17 | 1 | 0.03 |

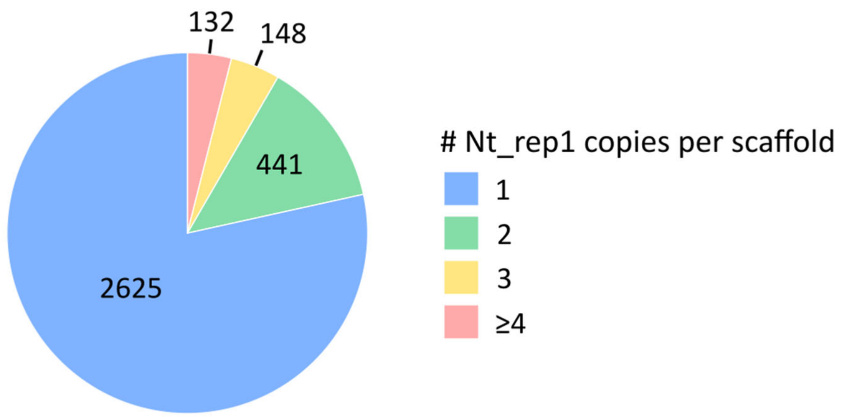

#### Figure S1.6.2.

Distribution of Nt_rep1 on the assembled scaffolds. Note that most of the scaffolds containing Nt_rep1 carry only a single copy.

Table S1.6.2. Overview of cytogenetic primers used in this study.

| Name | Sequence 5'-3' | Function | Reference |
|---|---|---|---|
| 18S-1 | CTG GTT GAT CCT GCC AGT AGT | 18S rDNA partial | (Jung and Lee 2012) |
| 18S-4 | GAT CCT TCT GCA GGT TCA CC | 18S rDNA partial | (Jung and Lee 2012) |
| "Ins_telo_F" | TAG GTT AGG TTA GGT TAG GT | Insect telomere motif | (Sahara, Marec, and Traut 1999) |
| "Ins_telo_R" | CTA ACC TAA CCT AAC CTA AC | Insect telomere motif | (Sahara, Marec, and Traut 1999) |
| Nt_pt1_F | ATG GTA TGG TAT GGT ATG GT | *N. tenuis* potential telomere motif 1 | This study |
| Nt_pt1_R | CAT ACC ATA CCA TAC CAT AC | *N. tenuis* potential telomere motif 1 | This study |
| Nt_pt2_F | TGG GTT GGG TTG GGT TGG GT | *N. tenuis* potential telomere motif 2 | This study |
| Nt_pt2_R | CCA ACC CAA CCC AAC CCA AC | *N. tenuis* potential telomere motif 2 | This study |
| Nt_pt3_F | CAG GTC AGG TCA GGT CAG GT | *N. tenuis* potential telomere motif 3 | This study |
| Nt_pt3_R | CTG ACC TGA CCT GAC CTG AC | *N. tenuis* potential telomere motif 3 | This study |
| Nt_rep1_F | TTC GCC CAA AAT GAA AAA ACG C | Nt_rep1 | This study |
| Nt_rep1_R | TCC TGA ACA AGT GTC TGT GTG T | Nt_rep1 | This study |

Figure S1.6.3.

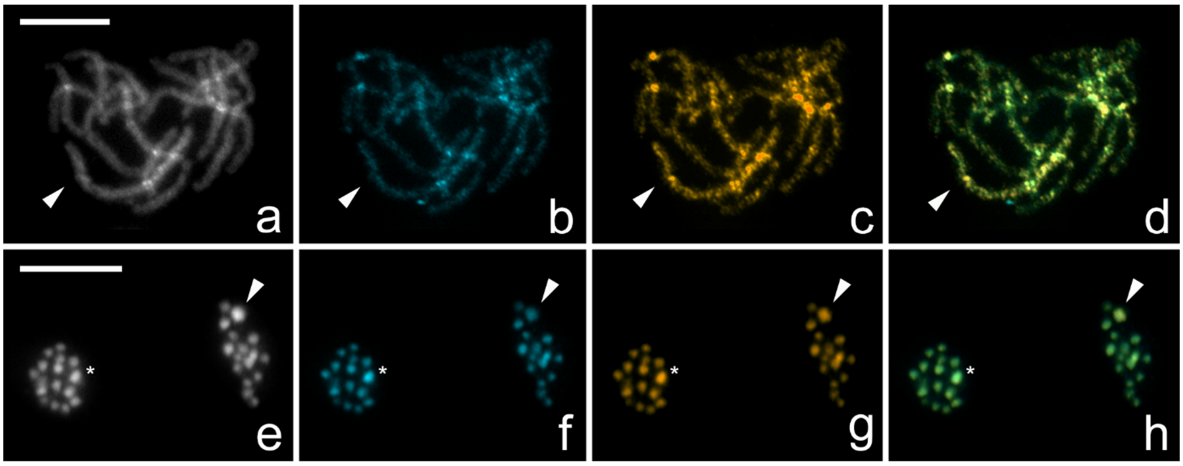

Alternate colouration for Figure 5 in-text. Comparative genomic hybridization (CGH) on female (**a**, **b**, **c**, **d**) and male meiotic metaphase II (**e**, **f**, **g**, **h**) chromosomes of *N. tenuis*. Panels (**a**, **e**) show chromosomes counterstained by DAPI (grey), panels (**b**, **f**) hybridization signals of the male derived genomic probe labelled by fluorescein (blue), panels (**c**, **g**) hybridization signals of the female derived genomic probe labelled by Cy3 (gold), and panels (**d**, **h**) merged images. (**c**, **d**) Note that the X chromosome bivalent (arrowhead) in female pachytene complement was highlighted more by female probe compared to the autosomal bivalents; (**b**, **d**) male probe labelled all chromosomes equally. (**h**) Two sister nuclei in meiotic metaphase II showed equal hybridization patterns of both probes on autosomes; in one of the forming nuclei, the X chromosome (arrowhead) was highlighted by female derived genomic probe (**g**, **h**) and in the second nucleus the Y chromosome (asterisk) was strongly highlighted by male derived genomic probe compared to autosomes (**f**, **h**) and less highlighted by female derived probe (**g**, **h**). (**e**) Note that the sex chromosomes are the biggest and most heterochromatic elements in the nucleus. Scale bar = 10 μm.

### S1.7. Poolseq results in full

Full results (*Tajima's D* and *Tajima's π*) of PoPoolation runs, under three conditions:

S1.9.1 Sliding window and step size 10K,

- original/S1.9.1 PoPoolationResults.csv

S1.9.2 Sliding window and step size 5K,

- original/S1.9.2 PoPoolationResults.csv

S1.9.3 Indel masking with sliding window and step size 10K (reported results).

- original/S1.9.3 PoPoolationResults_ReportedResults.csv

#### References

Jung, Sunghoon, and Seunghwan Lee. 2012. "Molecular Phylogeny of the Plant Bugs (Heteroptera: Miridae) and the Evolution of Feeding Habits." *Cladistics* 28 (1): 50–79. https://doi.org/10.1111/j.1096-0031.2011.00365.x.

Sahara, Ken, František Marec, and Walther Traut. 1999. "TTAGG Telomeric Repeats in Chromosomes of Some Insects and Other Arthropods." *Chromosome Research* 7 (6): 449–60. https://doi.org/10.1023/A:1009297729547.

Wheeler, David, Amanda J. Redding, and John H. Werren. 2013. "Characterization of an Ancient Lepidopteran Lateral Gene Transfer." *PLoS ONE* 8 (3). https://doi.org/10.1371/journal.pone.0059262.

Zhou, Weiguo, Francois Rousset, and Scott O Neill. 1998. "Phylogeny and PCR-Based Classification of Wolbachia Strains Using Wsp Gene Sequences." *Proceedings of the Royal Society B. Biological Sciences* 265: 509–15.
